# Modest changes in *Spi1* dosage reveal the potential for altered microglial function as seen in Alzheimer’s disease

**DOI:** 10.1101/2021.02.12.430903

**Authors:** Ruth E. Jones, Robert Andrews, Peter Holmans, Matthew Hill, Philip R. Taylor

**Affiliations:** Division of Infection and Immunity, Cardiff University, UK; UK Dementia Research Institute, Cardiff University, UK; Division of Psychological Medicine and Clinical Neurosciences, UK; Systems Immunity Research Institute, Cardiff University, UK

## Abstract

Genetic association studies have identified multiple variants at the *SPI1* locus that modify risk and age of onset for Alzheimer’s Disease (AD). Reports linking risk variants to gene expression suggest that variants denoting higher *SPI1* expression are likely to have an earlier AD onset, and several other AD risk genes contain PU.1 binding sites in the promoter region. Overall, this suggests altered levels of *SPi1* may alter microglial phenotype potentially impacting AD. This study determined how the mouse microglial transcriptome was altered following modest changes to *Spi1* expression in primary microglia. RNA-sequencing was performed on microglia with reduced or increased *Spi1*/PU.1 expression to provide an unbiased approach to determine transcriptomic changes affected by *Spi1*. In summary, a reduction in microglial *Spi1* resulted in the dysregulation of transcripts encoding proteins involved in DNA replication pathways while an increased *Spi1* results in an upregulation of genes associated with immune response pathways. Additionally, a subset of 194 *Spi1* dose-sensitive genes was identified and pathway analysis suggests that several innate immune and interferon response pathways are impacted by the concentration of *Spi1*. Together these results suggest *Spi1* levels can alter the microglial transcriptome and suggests interferon pathways may be altered in individuals with AD related *Spi1* risk SNPs.

## Introduction

Alzheimer’s Disease (AD) is the most prevalent form of Dementia, affecting millions of people worldwide [1]. Studies investigating AD genetics and pathology have suggested immune gene networks may contribute to an increased risk of developing AD [2,3]. *SPI1* encodes PU.1, a central transcription factor in microglial development and activation, and has a genome-wide significant genic association with AD in the IGAP GWAS (rs3740688 Odds Ratio 0.92 Meta p = 5.4 ×10^-13^), comprising 35,274 Alzheimer’s disease cases and 59,163 controls [4]. In addition, a 56 protein interaction network consisting of strongly enriched rare coding variants (p = 1.08 x10^-7^, and common variants with LOAD gene-wide significance (p = 2.98 x10^-7^) identified *SPI1* as a central hub gene [5]. Several studies have suggested *SPI1*/PU.1 levels impact on the microglial transcriptome, therefore impacting the phenotype of these cells. Several single-nucleotide polymorphisms (SNPs) associated with an increased risk of AD are thought to lie within the *Spi1* gene locus [3,6,7]. The SNP variant rs1057233^a^ is thought to result in a higher level of *Spi1* expression and earlier age of AD onset [7]. Moreover, *Spi1* is thought to influence the expression of other AD risk genes [5,7,8].

The PU.1 transcription factor is essential for the survival and function of macrophages [9–13] and is well conserved between humans and mice (>85% protein similarity, BLAST protein alignment (RefSeq ID’s NP_003111.2 and NP_035485, respectively). In hematopoietic development low levels of PU.1 drives B-lymphocyte development whereas cells expressing high levels of PU.1 are committed to the myeloid lineage [14–16]. In development PU.1 levels are regulated to commit precursors to a macrophage or B-cell lineage [17–23]. In these early experiments PU.1 appeared to have dose-dependent transcriptional thresholds in fetal liver macrophages [24]. PU.1 also regulates expression of several key macrophage receptors such as CSF1R, CD11b and CD45 [25–27]. Moreover PU.1 interacts with other lineage-determining factors such as C/EPBα/β to alter the chromatin landscape resulting in a specialised macrophage epigenome [28–30].

In both primary human microglia and the BV-2 mouse microglia cell line, reductions in PU.1 have resulted in changes to gene expression and a reduced phagocytic capacity [7,31,32]. Additionally increased PU.1 expression in BV-2 cells resulted in increased zymosan phagocytosis, and amplified both ROS, NO and cytokine production after LPS stimulation [32]. Though the impact of altered *Spi1* on the microglial transcriptome would ideally be studied in freshly isolated cells, such as those from a transgenic mouse model, at time of this study, no appropriate *Spi1* over-expression transgenic mouse lines were available.

CSF1R inhibitors are a potential AD therapeutic and prevent AD-associated microgliosis by blocking the CSF1R/PU.1 survival signalling pathway [33–36] but the impact on peripheral macrophages has not been reported. Knowing how subtle changes in *SPI1*/PU.1 levels contribute to the microglial tissue resident subset could allow more specific pathways could be targeted.

RNA-sequencing was used to assess changes to the microglial transcriptome following modest changes to *Spi1* expression in microglia from primary mixed-glia cultures. Modest changes to PU.1 protein were desirable as they reflect the expression changes caused by the risk allele, and therefore the biology underlying AD. A moderate reduction in microglial *Spi1* resulted in altered expression of cell cycle related genes while small *Spi1* increases upregulate immune response genes. A subset of genes identified as *Spi1* dose-sensitive highlighted a potential dose-dependent interferon driven immune response regulated by *Spi1*.

## Results

### RNA-Sequencing data shows impact of *Spi1* dose on microglia transcriptome

The effect of *Spi1* shRNA knock down (KD) and *Spi1* pSIEW over expression (OE) was assessed in flow cytometric sorted microglia after 11 days of lentivirus infection. Following *Spi1* knock-down *(Spi1* shRNA compared to non-silencing shRNA) 1615 genes were up- and 1462 down-regulated, using an adjusted p-value of <0.05 cut-off (Benjamini-Hochberg corrected for multiple testing). A proportion of down-regulated genes (230 genes) surpassed the −2 log2 fold-change cut-off (**Figure 1A**). In the *Spi1* overexpression dataset *(Spi1* pSIEW compared to pSIEW) 284 genes had an increased expression, 62 were down-regulated (p <0.05). In this comparison 71 up-regulated and 4 down-regulated exceeded a log2 foldchange value of 2. Principal component analysis (PCA, **Supplementary Figure 1**) confirmed that samples within the same group had greater similarity, and the *Spi1* pSIEW and *Spi1* shRNA samples clustered separately.

**Figure 1.**
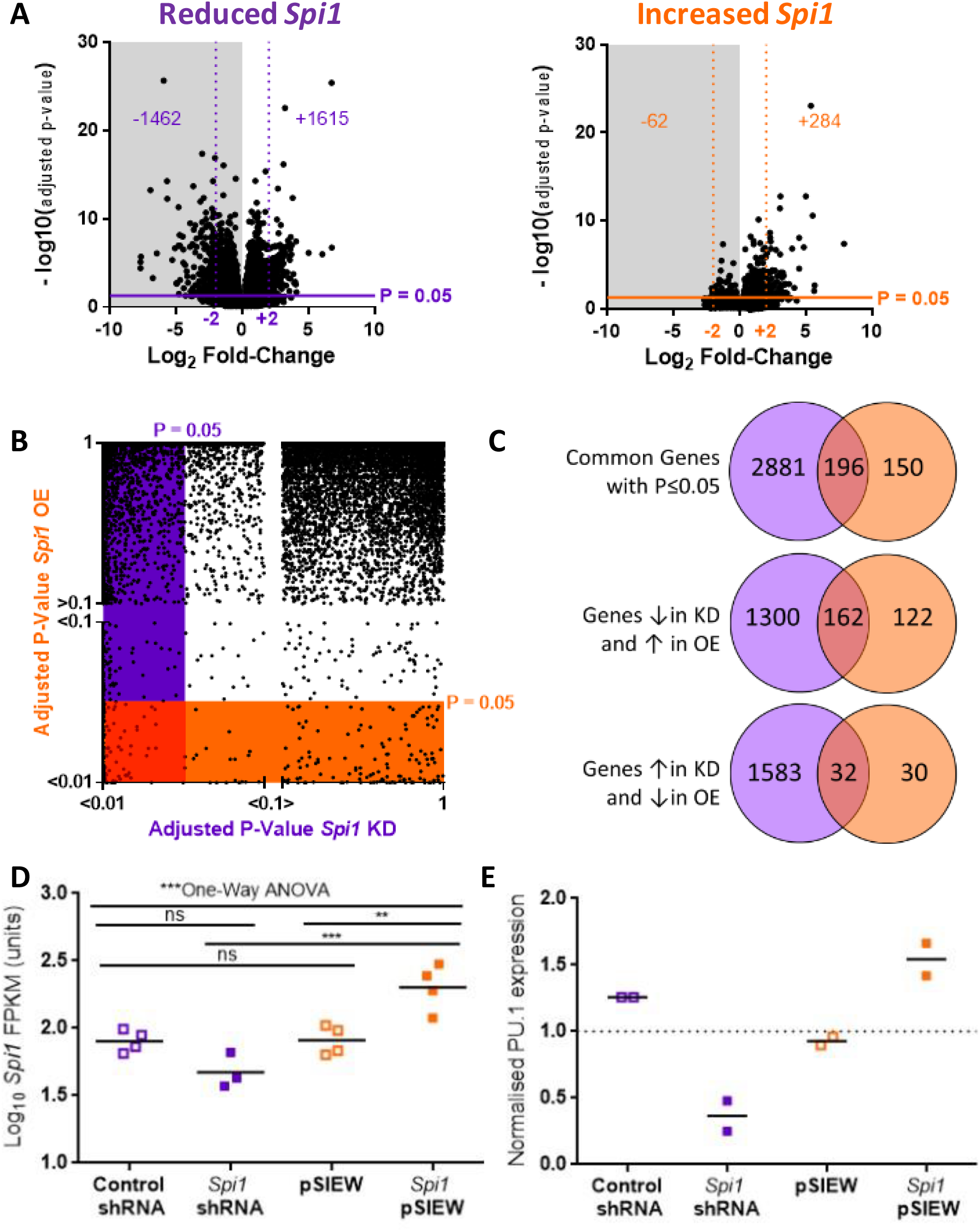
Results of RNA-Seq experiment in primary mixed-glial cultures. **A** Volcano plots summarising the distribution of genes in the Spi1 knock-down (purple) and Spi1 over-expression (orange). In the Spi1 knock-down dataset a 1462 genes were down-regulated (grey background) and 1615 up-regulated with a p-Value of ≤0.05 (solid lines), as indicated by the numbers on the graph. In the Spi1 over-expression dataset the majority of the genes, 284, were up-regulated and 62 genes had down-regulated expression. **B** Genes that were significantly changed in both the Spi1 knock down (KD, purple) and the Spi1 over expression (OE, orange) using a P≤0.05 threshold. A Plot of adjusted P-Values from all genes in both datasets. Genes that were below the P≤0.05 threshold in the Spi1 knockdown dataset are highlighted in purple, those that were below the P≤0.05 cut-off in the Spi1 over-expression are within the orange bar. In the bottom left corner the red line surrounds the 196 genes that were significantly changed in both datasets. **C** Venn diagrams summarising the gene expression that were significantly changed (P ≤ 0.05) in either the Spi1 knock-down dataset (purple), the Spi1 over-expression dataset (orange) or changed in both datasets (red). Of interest were the genes with expression that appears to be sensitive to the Spi1 dose in microglia, namely the 194 genes that appear to be expressed relative to the dose of Spi1. **D** The logw number of Spi1 mRNA fragments per Kilobase of transcript per Million mapped reads (FPKM) in the RNA-sequencing datasets. Each dot represents the value from one biological replicate, and the means is indicated by the horizontal lines. One-way ANOVA on the logw transformed data confirmed significant differences between the group means (***p=0.0003). Spi1 mRNA expression was increased in the samples infected with the Spi1 over-expression dataset (filled orange) compared to empty vector control samples (outline orange) (~2.5x increase, Sidak’s multiple comparison test **p≤0.01). FPKM values for Spi1 shRNA samples did not significantly differ from control samples (Sidak’s multiple comparison test p>0.05). Spi1 pSIEW and Spi1 shRNA groups significantly differed whereas control samples did not (Sidak’s multiple comparison test ***p≤0.001 andp>0.05 respectively). **E** PU.1 protein expression in primary microglia cultures treated with lentiviruses designed to manipulate Spi1 mRNA expression. PU.1 protein expression was normalised to the non-infected (NI) samples in each experiment to provide a consistent baseline (as described in methods). The bar denotes the mean and the individual replicates by symbols. Microglia infected with the Spi1 shRNA (solid purple) have approximately a 60 % reduction in PU.1 expression compared to cells infected with a non-silencing shRNA control (lined purple). The Spi1 over-expression virus (Spi1 pSIEW, solid orange) increased PU.1 protein expression in microglia by roughly half compared to cells infected with an empty vector control (lined orange). Both experiments were performed with mixed glia cultures from 8-week-old female C57BL/6J mice.

The datasets with altered *Spi1* expression were then compared to identify which subset of genes were likely affected by *Spi1*/PU.1 in a dose-dependent manner. This identified 196 genes which were significantly diminished in the *Spi1* knock-down and upregulated in the *Spi1* over-expression dataset (Adjusted P-Value of ≤ 0.05; **Figure 1B**). When the direction of the gene fold-changes were compared 162 of these genes were lower in the dataset with lower *Spi1* and higher in the *Spi1* over-expression dataset and 32 genes where expression was increased following *Spi1* knock-down and reduced in the *Spi1* overexpression dataset (**Figure 1C**). Therefore, 194 genes were classed as *Spi1* dose-sensitive (**Supplementary Table 1**).

RNA-sequencing confirmed *Spi1* mRNA expression was increased in the *Spi1* pSIEW samples (One-Way ANOVA on Log10 data p=0.0003, Sidak’s multiple comparison p≤0.01) though no significantly difference was determined at the mRNA level between *Spi1* shRNA replicates and the appropriate control viruses (Sidak’s multiple comparison test p>0.05, **Figure 1D**), although shRNA can significantly affect translation with minimal impact on mRNA levels.

PU.1 protein was assessed in independent microglia samples (**Figure 1E**) and appeared to be reduced by ~60 % in Spi1 shRNA samples compared to control shRNA infected microglia. *Spi1* over-expression in pSIEW lentivirus infected microglia increased PU.1 protein expression by ~50 % in vitro microglia compared to uninfected cells and to microglia infected with pSIEW control. The values clearly cluster within each biological group and the means of *Spi1* shRNA and *Spi1* pSIEW microglia are disparate it suggests PU.1 was altered by *Spi1* modulating viruses.

The differential affect of the viruses on mRNA and protein levels suggests that the *Spi1* shRNA may block translation rather than degrading *Spi1* mRNA to reduce PU.1 protein expression.

### Gene Ontology analysis in *Spi1* KD and *Spi1* OE datasets

Differentially expressed genes (p <0.05 significance threshold) in the *Spi1* knock-down and *Spi1* overexpression datasets were separately assessed using DAVID (Database for Annotation, Visualization and Integrated Discovery, version 6.8). **Figure 2** displays the Gene Ontology (GO) terms from 20 most significantly enriched pathways in each dataset (corrected p values p < 0.05, FDR). Primary microglia with a lower *Spi1* expression the most significantly changed pathways were related to cell cycle and DNA repair (**Figure 2A**). However, microglia with higher *Spi1* (**Figure 2B**) had an over representation of genes related to antigen presentation pathways, immune system processes and response to interferon.

**Figure 2.**
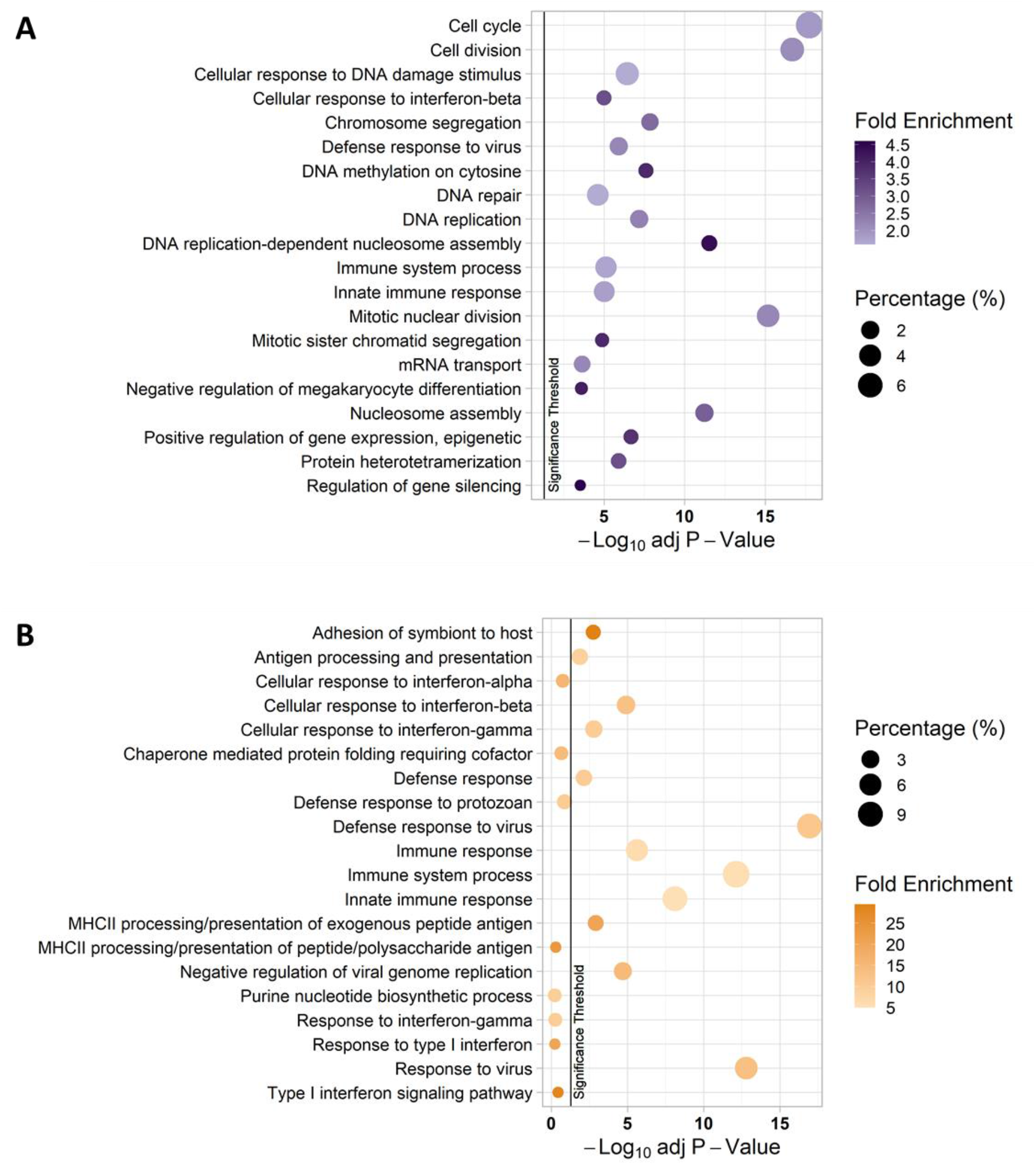
The 20 most significantly changed biological pathways in the Spi1 knock-down (A) and Spi1 over-expression (B) datasets assessed using DAVID. In these graphs the Benjamini-Hochberg adjusted P-value(-Log10) is displayed on the x-axis, the Gene Ontology (GO) term listed on the y-axis, the percentage of the gene list in each cluster is denoted by the size of the bubble and the colour denotes the fold-change, where a darker colour indicates a stronger enrichment. The vertical black line indicates the p-value cut-off of 0.05.

### *Spi1* Dose-Sensitive Subset highlight interferon response pathways

The absolute log2 fold-changes of 162 genes reduced by *Spi1* knock-down and increased by *Spi1* overexpression significantly correlated to each other (**Figure 3A**) and the 32 genes that were increased by *Spi1* knock-down and reduced by *Spi1* over-expression (**Figure 3B**) were also found to correlate (Two-tailed Spearman Rank Test approximate p<0.0001 and r >0.8 for both). This further supports the hypothesis that the expression of these genes is controlled in a *Spi1* dose-sensitive manner. Collective analysis of the 194 *Spi1* dose-sensitive gene list using GO terms through DAVID indicated an enrichment of immune response genes (**Figure 3C**), particularly pathways linked to interferon and viral defence responses. These results suggest that a high microglial *Spi1* expression may result in more responsive microglia that produce more interferon, though further work would be required to confirm this experimentally.

**Figure 3.**
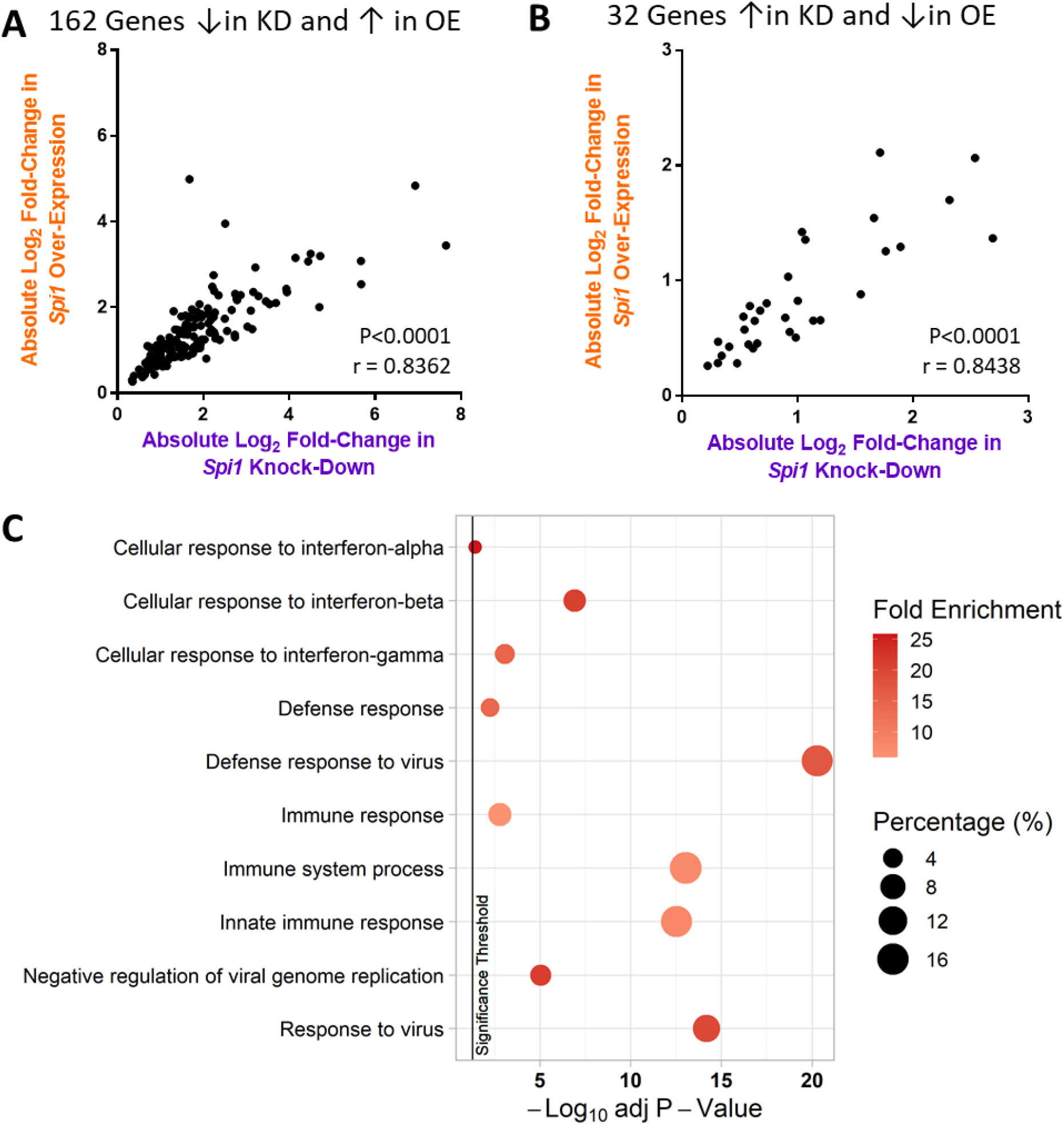
Absolute Log2 fold-changes in Spi1 dose-sensitive genes. **A** Absolute log2 fold-change values were used the 162 genes where p≤0.05 with negative fold-change values in the Spi1 knock-down dataset (purple) and positive foldchange values in the Spi1 over-expression dataset (orange) were assessed separately from the 32 genes where p≤0.05 and Spi1 dose had the opposite effect on expression (**B**). Two-tailed Spearman Rank test identified significant correlations between the log2 fold-changes in both graphs (approximate p<0.0001 and r > 0.8). **C** Top 10 most significantly altered “Biological Processes” according to the Benjamini-Hochberg adjusted P-value. The bubble size indicates the percentage of the gene list aligned to this pathway. The colour indicates the fold enrichment, which is the proportion of genes present in this list compared to the background gene expression. The vertical black line indicates the p-value threshold of 0.05.

### Several Gene Clusters Implicate Immune System Responses

The data were further assessed using hierarchical clustering of the FPKM expression values (**Figure 4**). Six discrete clusters were identified, and gene lists were assessed using DAVID as before. Genes with an increased expression in microglia overexpressing *Spi1* (Cluster 1) are enriched for pathways involving MHCII processing and presentation of antigens. The genes in Cluster 2 appear to be the most *Spi1* sensitive, where expression was reduced in the *Spi1* shRNA and increased in the *Spi1* over-expression samples compared to the respective controls. Most pathways associated with the genes in this cluster are related to the immune response pathways including viral response pathways. Cluster 3 contained genes that were down-regulated in the *Spi1* shRNA samples compared to the controls and contains genes related to interferon signalling and immune response pathways. Cluster 5 contains genes that had a higher expression in the *Spi1* shRNA samples compared to the control shRNA control samples. The GO TERMS associated with the genes in this cluster include those related to cell cycle and DNA replication pathways. Overall, this reinforces previous reports that reduced *Spi1* seems to impact cell cycle which was not unexpected given PU.1’s involvement in survival signalling [11,37]. A lowering of *Spi1* appeared to reduce activation of immune response related pathways and vice versa in the samples where *Spi1* was overexpressed.

**Figure 4.**
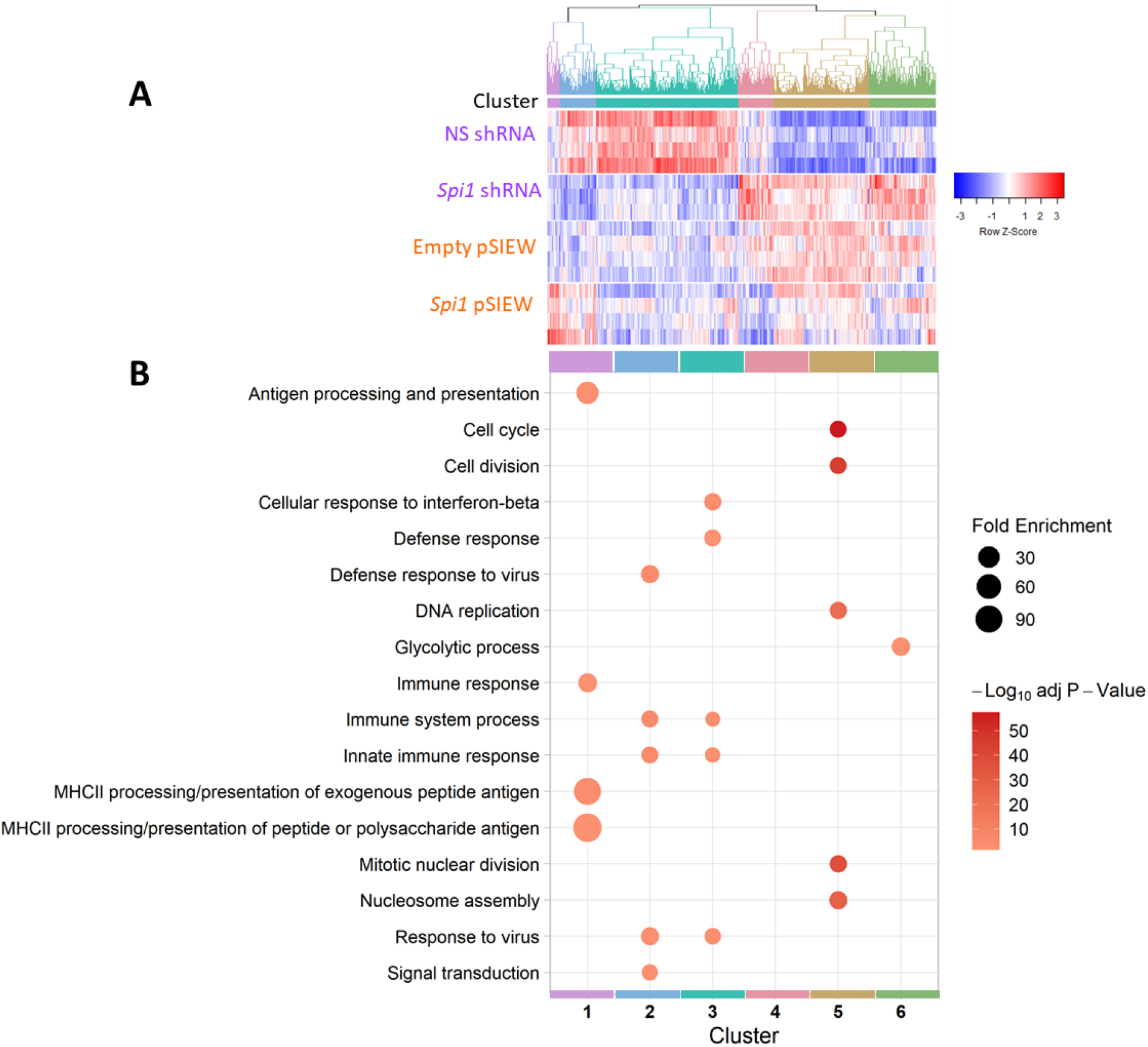
Hierarchical clustering analyses with the 20 most siginificantly changed biological pathways each cluster assessed using DAVID. **A** Hierarchical clustering of rows, where each row represents the scaled log10 FPKM values for each gene, with dendrogram higlighting cluster boundaries. The Pearson correlation was used to calculate the Z-Scores and UPGMA agglomeration methods were used. These 6 clusters were produced using the cutree function at a height of 1.59. **B** The gene lists were assessed using DAVID and a bubble plot created to highlight the top 5 most significatly enriched pathways (P-Value threshold of <0.05) in each gene cluster. Cluster 1 contains genes that had increased expression following a higher Spi1 expression and are mainly linked to immune responses and MHCII antigen presentation. Cluster 2 appears to contain more of the Spi1 dose-sensitive genes and are enriched for viral/immune response signalling pathways. Genes in Cluster 3 had a lower expression in the Spi1 shRNA dataset and are related to immune response and interferon signalling pathways. Cluster 4 contained no significantly enriched pathways using the p≤0.05 threshold. Cluster 5 contains multiple genes related to cell cycle and DNA replication pathways, whose expression was increased in the Spi1 shRNA samples. The bubble size denotes the fold enrichment and the colour Benjamini-Hochberg adjusted P-values.

### PU.1 dose-related transcriptomic differences were not limited by direct binding

Changes in gene expression after manipulation of PU.1 could arise from changes in direct DNA binding or via secondary effects. To identify possible direct targets of PU.1 that also had altered expression, the proportion of protein coding genes that contained a PU.1 binding site sequence or promoter region were assessed within each dataset and the subset of *Spi1* dose-sensitive genes utilising a published ChIP-Sequencing (ChIP-Seq) dataset available online [30]. **Figure 5** shows ~70 % of genes differentially expressed in the RNA-Seq datasets contained a PU.1 binding site and therefore expression was potentially directly modified by *Spi1.* Moreover, 30-40 % of genes in *Spi1* KD and *Spi1* OE datasets had PU.1 binding sites within the promoter region suggesting their expression was directly altered by *Spi1*/PU.1. These proportions remained consistent even in the *Spi1* dose-sensitive subset. Only differentially expressed genes in the *Spi1* KD were enriched for PU.1 binding sites in the promoter region (Fisher’s exact test, p-value of overlap=1.6×10^-16^). Given the proportion of overlapping genes was similar across all three test sets *(Spi1* KD, *Spi1* OE and *Spi1* dose-sensitive) it seems *Spi1* function was not limited to genes it directly regulates, but likely indirectly mediates expression of multiple other genes via secondary downstream processes.

**Figure 5.**
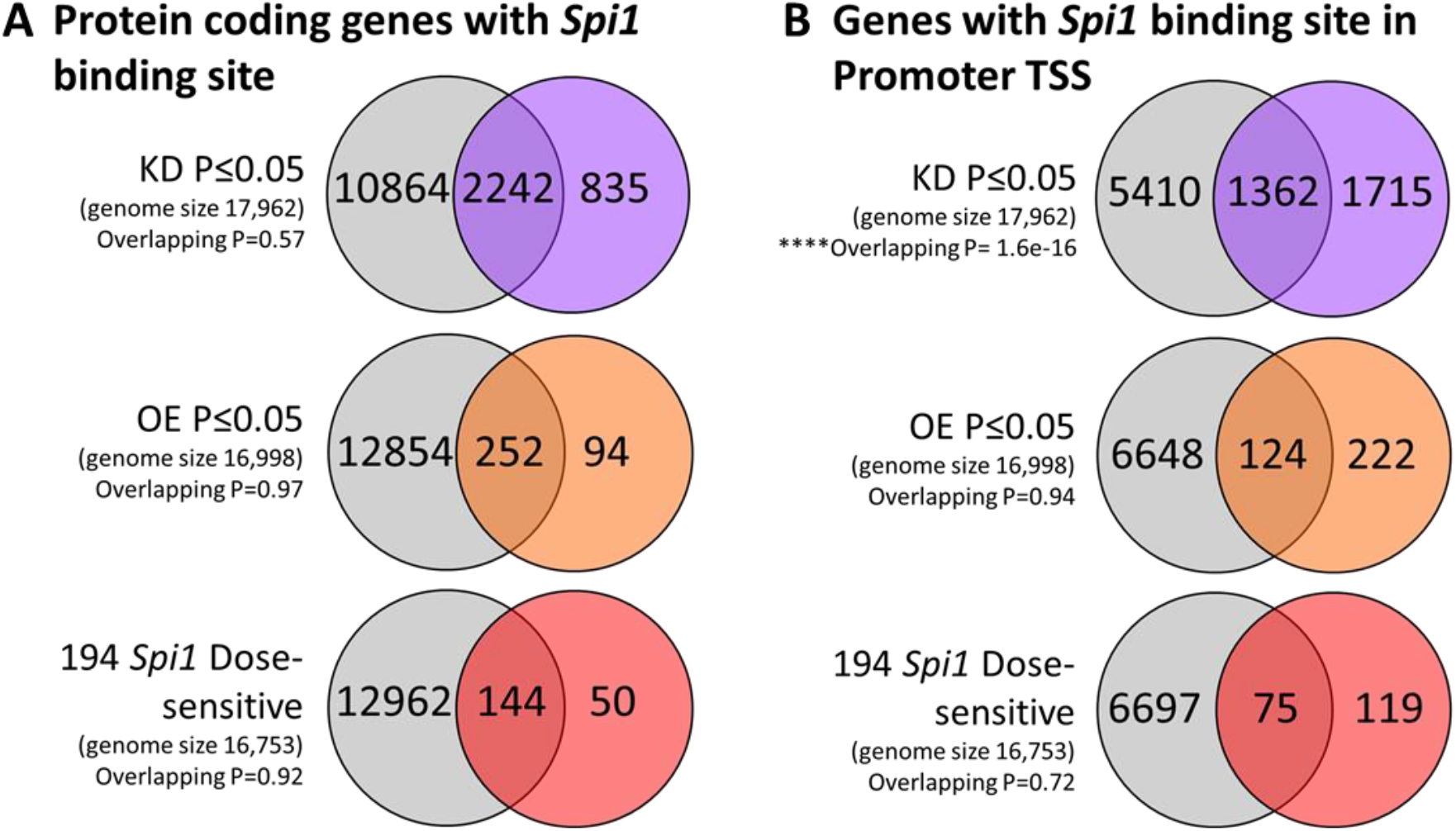
Comparison of Spi1 KD, Spi1 OE and 194 Spi1 Dose-sensitive genes to Spi1/PU.1 Chip-Seq dataset. Spi1 binding sites were determined from PU.1 Chip-Seq published data available online [30] as described in methods. Fisher’s exact tests were used to determine if there was a significant overlap between the datasets **A** includes the results from all protein coding genes with a Spi1 binding region (defined by Chip-seq in grey) that are also expressed in KD (purple), OE (orange) RNA-seq datasets or the Spi1 dose-sensitive genes (red). **B** The proportion of genes expressed in the RNA-seq datasets that contained a Spi1 binding site within the promoter region. The KD dataset contained a significant number of genes with a Spi1 binding sequence in the promoter (Fisher’s exact test, overlapping p-value = 1.6×10^-16^). No significant enrichment was noted in either the Spi1 OE or dose sensitive genes.

### Comparison to human AD risk genes

Protein network analysis suggest *Spi1*/PU.1 is one of several “hub” genes within the network of AD risk genes [5], which was further supported by Cis-eQTL analyses in monocytes and macrophages [7]. The RNA-Sequencing datasets generated in this paper were compared to the International Genomics of Alzheimer’s Project (IGAP) dataset, to investigate if there was an enrichment of AD genetic risk within the differential expressed sets of genes. **Supplementary Table 2** denotes the gene sets used in this MAGMA analysis. In the *Spi1* over-expression dataset a 21 gene set (**Figure 6**), corresponding to a Benjamini-Hochberg adjusted P-value of ≤1×10^-6^ for differential expression, were enriched for AD genetic risk (adjusted p-value = 0.035). The individual IGAP p-values can be viewed alongside the *Spi1* over-expression dataset values in **Supplementary Table 3**. This gene list (**Figure 6B**) contained *Ifit3, Oas1b* and *Oas2* which GO analysis linked to the interferon response pathway and *Rnf144b* and *Treml4* which are related to antigen presentation pathways, implicating the immune system response in AD pathology.

**Figure 6.**
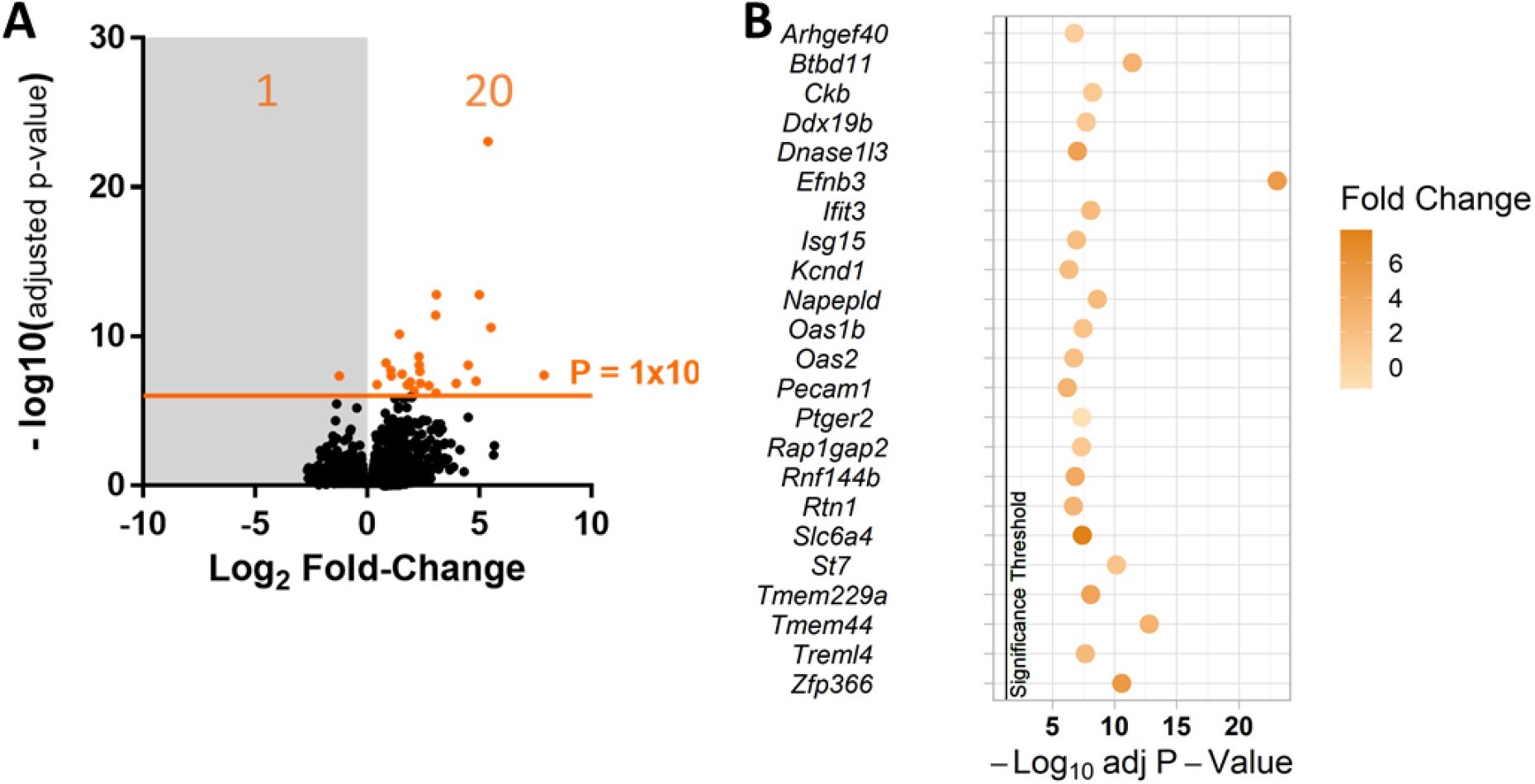
Summary of Spi1 over-expression genes that were significantly enriched in the IGAP dataset. **A** Volcano plot of the Spi1 over-expression gene expression changes relative to the empty vector control, highlighting the p ≤1×10^-6^ threshold that was shown to be significantly enriched by MAGMA analysis in the IGAP dataset (MAGMA’s empirical multiple testing corrected p-value 0.035). The adjusted P-value for this gene set was 0.0033 and an enrichment effect size of 0.51. **B** Bubble plot of the genes from the Spi1 over expression dataset that were enriched for AD genetic risk via MAGMA comparison to the human IGAP database, against the Benjamini-Hochberg adjusted P-Values from the Spi1 over-expression RNA-sequencing data with the fold-change shown in colour. The vertical black line indicates the P-value threshold of 0.05.

## Discussion

Manipulation of *Spi1* expression in primary mixed glia cultures were used to identify how microglial gene expression was altered following modest changes to *Spi1*/PU.1. Analysis of the RNA-sequencing datasets produced from these cultures have provided insight into how *Spi1* dose affects the microglia transcriptome. Briefly *Spi1* knock-down resulted in changes to gene expression of proteins enriched for cell cycle checkpoint pathways and over-expression of *Spi1* altered enriched pathways related to MHCII, interferon response and viral response (**Figure 3** & **Figure 4**). Comparing these RNA-Seq datasets to ChiP-Seq data suggests that approximately 70 % of the differentially expressed genes were likely directly altered by *Spi1* binding whereas the rest altered as a downstream consequence following changes to other genes (**Figure 5**).

Strengths of this experimental approach include assessing the bidirectional impact of *Spi1* changes on the microglial transcriptome in an unbiased manner in primary cell cultures. Whilst we supplemented our cultures with TGF-β, which partially compensates for the loss of complex environmental cues microglia would normally receive, it does not fully replicate the *in vivo* context and there were no suitable transgenic mice available to study the impact of *Spi1* dose on microglia *in situ.*

PU.1 has been previously linked to proliferation in macrophages. Bone-Marrow Derived Macrophages over-expressing PU.1 increased GM-CSF and M-CSF dependent proliferation and cell number, the opposite was seen in cells transfected with an anti-sense PU.1 [9].

However, there are some conflicting reports in the literature as to whether PU.1 alters microglia proliferation. In this report absolute cell numbers did not differ between *Spi1* shRNA and control shRNA infected microglia, neither were there significant fold-change differences between normalised cell cycle data (**Supplementary Figure 2**). Given that in human microglia cultures an siRNA mediated reduction of PU.1 resulted in a lower cell number, disintegration and rounding of some microglia was observed but at the final timepoint viability appeared unaffected suggesting some cells could survive with reduced PU.1 [37]. Another similar study saw no reduction in microglia number, though PU.1 loss was measured at a culture level and the impact of a ~60% reduction on individual cells not observed [31]. In summary, whilst we did not observe significant changes in cell cycle associated with limited changes in PU.1 levels, we did observe changes in cell cycle associated genes with such PU.1 level changes.

As PU.1 has been shown to bind the promoter of the Csf1-receptor (Csf1r) increasing expression of Csf1-receptor [38], which is highly linked to microglia survival and proliferation. In BV-2 cells a reduction in PU.1 also results in a reduced *Csf1r* expression in BV-2 cells [7] though no reports were given on cell number/viability. It was recently shown that BV-2 cells with diminished PU.1 expression were more vulnerable to caspase-dependent cell death, while PU.1 over-expressing cells appear to have delayed onset of death, though baseline cell viability did not appear to be impacted by PU.1 modulation [32]. In the *Spi1* RNA-seq datasets generated in this paper *Csf1r* expression was not significantly altered in either the knock-down or over-expression datasets (Benjamini-Hochberg adjusted P-Values 0.665 and 0.999 respectively), though the impact on Csf1r protein expression was not measured in these samples. Given that Csf1r inhibitors have been shown to prevent AD-related microgliosis and partially ameliorate disease pathology *in vivo* [33–36] it would be interesting to see if Csf1R protein was affected in *Spi1* shRNA infected microglia and if the reduction in PU.1 had a similar impact to Csf1r inhibitors on disease progress.

Over-expression of *Spi1* resulted in changes to gene expression of proteins enriched for MHCII related pathways, interferon response pathways and response to virus pathways (**Figure 2B**). Though there may be minor concerns as to the significance of viral immune response pathways could be an unintended effect of using lentiviruses to manipulate *Spi1* expression this is unlikely as comparison were made relative to a control virus which was identical bar the *Spi1* coding sequence. Additionally, this study identified a subset of 194 genes that appear to be *Spi1* dose sensitive, enriched for GO terms such as “Cellular response to interferon” and “Defence response to virus” (**Figure 3**). Therefore, an increased level of *Spi1* appears to cause transcription of genes associated with an inflammatory phenotype that might be relevant to AD genetic risk mechanisms (**Figure 6**).

Previous work has identified that *Spi1* is able to influence multiple gene networks in microglia [5,7,31] and has been proposed to be central to a network of AD risk genes that are conserved between species [39]. Basal cytokine expression was higher in BV-2 cells over-expressing PU.1, and was further increased by LPS stimulation compared to control samples [32]. While media taken from PU.1 over-expressing LPS treated human induced microglia-like cells (iMGLs) promoted a reactive astrocyte phenotype in primary cultures this effect was not seen in astrocytes given media from LPS treated iMGLs with reduced PU.1 expression [32]. In co-transfected NIH-3T3 fibroblasts PU.1 binding to either IFN Regulator Factor 4 (IRF-4) protein or IFN Consensus Binding Protein (ICSBP) was required for maximal induction of an IL-1β luciferase reporter [40]. Therefore, it is likely that increased *Spi1*/PU.1 expression levels in microglia may result in a more inflammatory microglial profile, potentially in conjunction with these additional transcription factors. Gene ontology analysis of the *Spi1* knock-down and over-expression datasets suggest that the level of *Spi1* may impact microglial phenotype. While others have confirmed PU.1 dose changes result in phenotypic differences in BV-2 cells [32] further profiling is needed to ensure the pathway analysis matches the phenotype presented in primary microglia or ideally *in vivo.*

Previous reports also suggest a potential link between *Spi1* binding sites near other AD risk genes or regulatory elements [5,7,8]. *Spi1* risk alleles associated with a higher gene expression have been shown to lower the age of onset in AD, highlighting the need to understand how *Spi1* contributes to microglia phenotype [7]. Work in BV-2 cells suggests that reduced PU.1 results in a less reactive phenotype whereas PU.1 over expression primes cells for a more reactive response [32]. Salih *et al.* (2019) identified a network of microglial genes expressed by amyloid-associated microglia, collated from five transgenic mouse lines, included *Spi1,* and 74 other genes that were identified as *Spi1* dose-sensitive in this study [41]. These genes included *Oas1b, Oas2, Ifit3* which were all highly differentially expressed in the *Spi1* over-expression dataset (**Figure 6**) and are linked to interferon signalling [42]. Moreover, Sierskma *et al.* (2020) identified 18 genes common in APP transgenic mouse model over age and AD GWAS datasets, which were predominantly expressed by microglia and appear to be regulated by *Spi1.*

In summary, disease relevant reductions in *Spi1* highlighted dysregulation of genes involved several cell cycle checkpoint pathways. Modest increases in microglial *Spi1* expression results in dysregulation of genes linked to immune response and interferon signalling, suggestive of a more pro-inflammatory microglial phenotype. This study highlights how relatively modest changes to *Spi1* expression alone can have a large impact on the microglial transcriptome of primary mixed-glial cultures, providing candidate pathways for future studies investigating *Spi1* dependent processes and AD relevant biology.

## Methods

### Primary Mixed Glia Cultures

Brains from 8-week-old C57BL/6J mice (Charles River) were collected and transported in Hank’s Balanced Salt Solution (HBSS without Mg^2+^ or Ca^2+^; Gibco) on ice. Three brains were used per experiment which was repeated times 7 times (n=7), 4 to generate RNA-Sequencing replicates and 3 for subsequent validation experiments (**Supplementary Figure 3**). Brains were digested using the Neural Tissue Digest Kit-P in C-tubes as per manufactures directions (Miltenyi Biotec). Two brains were digested in each C-tube using program 37_ABDK on the GentleMACS™ OctoDissociator (Miltenyi Biotec) to produce a single cell suspension. Cell suspensions were passed through a 70 μM strainer (Falcon) into a 50 mL tube and centrifuged at 300 *x g* for 7 minutes. The supernatant was aspirated and the pellet resuspended in 10 mL Dulbecco’s Minimum Essential Media (DMEM containing 4.5 g/L D-Glucose, GlutaMAX™ and supplemented with 15 % (v/v) heat-inactivated Foetal Bovine Serum (FBS) and 100 units/mL (v/v) Penicillin and 100 μg/mL Streptomycin (v/v); all Gibco) before centrifugation at 300 *x g* for 7 minutes. The supernatant was aspirated, and the pellet resuspended in 12 mL of DMEM media containing 10 ng/mL recombinant murine M-CSF (Peprotech). Cell suspensions were then pooled as required and placed in a 100 mm x 20 mm (diameter x height) tissue culture plates, 12 mL per plate, and moved to a humid incubator with 95 % air and 5 % CO2 overnight to allow the microglia to adhere.

Myelin was resuspended in the media (with 24 hours of digest) by gentle plate agitation and the media containing the myelin debris was discarded. The cell monolayer was carefully washed with 10 mL of DMEM media. The wash media was then replaced with 12 mL DMEM media with 10 ng/mL M-CSF and this was again replaced two days later. On day 5 of culture the media was replaced with DMEM media containing 10 ng/mL M-CSF and 50 ng/mL recombinant murine TGF-β1 (eBioscience™). TGF-β supplementation was used to promote a more homeostatic phenotype *in vitro* [43,44]. From this point forward all culture media contained M-CSF and TGF-β1 was replaced every two days. On day 10, these cultures were infected with *Spi1* shRNA, control shRNA, *Spi1* pSIEW or pSIEW lentivirus particles as appropriate. To achieve this the culture media was replaced with 3 mL fresh media and 300 μL of lentivirus was added to each plate. After 6 hours each plate received an additional 3 mL of culture media. The media on these cultures continued to be changed every 2 days, supplemented with M-CSF and TGF-β as before.

### Sample Processing for RNA-Sequencing

After 21 days the media was removed, set-aside, and the mixed glial cells harvested by incubating ~10 mL Accumax™ (Sigma) at 37 °C for 5 minutes. The monolayer was gently washed with DMEM media and any remaining attached cells were carefully removed using a plastic scraper (Greiner). The cell suspension was added back to the media and centrifuged at 300 *x g* for 7 minutes. The supernatant was removed via pipetting and the cell pellet was resuspended in 0.5 % BSA in DPBS with a 1:1000 dilution of LIVE/DEAD® near-IR staining solution for 30 minutes on ice per manufacturers direction (Molecular Probes).

Samples were centrifuged at 300 x g for 7 minutes and the supernatant aspirated via pipette. Each sample was resuspended in 500 μL block solution (4 μg/mL Rat anti-mouse FcγRII/III (clone 2.4G2) and 5 % (v/v) filtered rabbit serum in 0.5 % BSA (w/v), 5 mM EDTA in DPBS) and kept on ice for 10 minutes. CD11b (Clone M1/70 PerCP-Cy5.5 conjugate, BD Biosciences final dilution 2 μg/mL) and CD45 (Clone 30-F11 eFlour® 450 conjugate, eBioscience™ final dilution 2 μg/mL) antibodies were diluted to a 4 μg/mL concentration in 0.5 % BSA (w/v), 5 mM EDTA in DPBS. 500 μL of the antibody staining solution was added to each sample, to give a final antibody concentration of 2 μg/mL, and the samples were incubated on ice for a further 15 minutes. The tubes were then centrifuged at 300 *x g* for 7 minutes and the supernatant removed.

The cells were resuspended in 1 mL of 0.5 % BSA (w/v) and 5 mM EDTA in DPBS and kept cool before undergoing fluorescent activated cell sorting on a FACS Aria™ III (BD Biosciences). Dead cells were excluded from the sort and microglia were selected using CD11b/CD45 double staining and GFP as a marker of infection. The sorted cells were then pelleted via centrifugation, the supernatant aspirated and the pellet lysed for RNA using the Mini RNeasy kit (Qiagen) per manufacturer’s directions.

### Lentivirus Preparation

The shRNA vector construct and lentivirus production method has been described in [45] and sucrose-gradient purification in [46]. The insert sequences for the non-silencing shRNA and *Spi1* shRNA can be seen in **Table 1**.

**Table 1.**
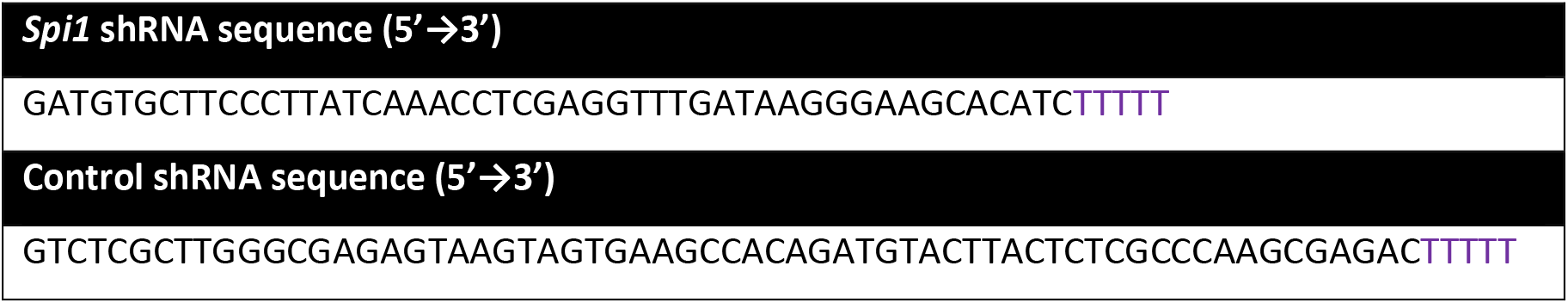
**shRNA sequences** including termination sequence (purple).

Briefly the over-expression plasmid used a spleen focus forming virus (SFFV) promoter to drive expression of murine *Spi1* (Ensemble reference CCDS 16425.1). An Internal Ribosome Entry Site (IRES) was used to initiate translation of a downstream eGFP reporter. Lentiviruses were purified by overlaying the virus containing media over a 20% sucrose solution before ultracentrifugation at 26,000 rpm for 90 minutes at 4 °C in a SW28Ti swinging-bucket ultracentrifuge rotor assembly in an Optima XPN-80 Ultracentrifuge (Beckman Optima Ultracentrifuge) and the viral pellet was resuspended in AimV™ media.

### RNA-Sequencing Data Anaysis

RNA was isolated from cell sorted GFP+ microglia using the RNeasy Mini kit (Qiagen) per manufacturer’s instructions and eluted in nuclease free water. RNA integrity and concentration were assessed using the Agilent 2100 Bioanalyzer (Aligent). Complementary (cDNA) libraries were generated using the Truseq® stranded total RNA with Ribo-Zero GOLD kit (Illumina®). Paired end sequencing was then performed using the Illumina® HiSeq 2500 sequencing platform to a read depth of between 30-40 million pairs

The FASTQ files were processed with Trimmomatic [47] to remove paired-end reads and quality was confirmed in FastQC using default parameters [48]. Following this sequence reads were mapped to the mm10 genome (GRCm38) using the STAR pipeline [49] and featureCounts was used to assign counts to transcripts [50] with the GRCm38.84 Ensembl gene build GTF. The Ensembl FTP site was used to download the reference genome and GTF [51]. Differential gene expression was performed using the DESeq2 package [52]. Genes that were not significant after the differential expression analysis were discarded; significance was defined as an adjusted P-value (Padj) of <0.05 (Benjamini-Hochberg correction for multiple testing).

#### Assessing PU.1 promoter binding sites

Briefly, a microglial PU.1 ChIP-sequencing (ChIP-seq) dataset (GSM1533906) isolated from C57BL/6 mice of a similar age (8-9 weeks) [30] was accessed via the Cistrome database [53]. This dataset was then run through HOMER [28] to generate an annotated peak file with genomic features. The promoter region was defined as 1000 base pairs upstream or 100 base pairs downstream of the transcription start site.

Duplicates were removed from the annotated ChIP-seq dataset and compared to gene discoveries from knock-down, over-expression and dose-sensitive *Spi1* RNA-sequencing datasets. The background genome size was the number of genes detected in each RNA-Sequencing dataset defined as either all control and/or all experimental samples had a raw read count of ≥5. In the *Spi1* knock-down dataset 17,962 genes were detected and 16,998 genes in the *Spi1* over-expression data. For the 194 *Spi1* dose-sensitive genes a merged list of genes expressed in either the *Spi1* knock-down or over-expression RNA-sequencing datasets was generated, and duplicates removed, to give a background gene number of 16,753.

Finally, the ChIP-seq datasets were compared against their respective RNA-Seq dataset (Padj ≤0.05) in R using the ‘TidyVerse’ and ‘GeneOverlap’ packages. Here, a Fisher’s exact test utilised to confirm if there was any significant enrichment of PU.1 ChIP-seq binding sites in genes expressed in *Spi1* knock-down, over-expression or dose-sensitive datasets.

#### Gene Ontology Analysis

Enriched gene Ontology (GO) terms were identified using the Database for Annotation, Visualization and Integrated Discovery (DAVID, version 6.8) and compared against a background of genes expressed in at least one *Spi1* RNA-seq dataset (16,998 genes). Gene expression in the *Spi1* RNA-seq datasets was defined as having a raw read count ≥5 in all control and/or all experimental samples. The most significantly altered terms from the GOTERM_BP_DIRECT list, henceforth called Biological Process GO Terms, were downloaded for subsequent analysis. Bubble plots were generated in R using the ‘ggplot2’ package, and an adjusted P-value cut-off of 0.05 indicated on each plot. Data utilised in these plots included the Benjamini-Hochberg adjusted p-value, the percentage of the genes from the RNA-seq datasets that aligned to this pathway and the fold enrichment, which was defined as the proportion of genes present in this list compared to the *Spi1* RNA-Seq gene expression background.

#### Hierarchical clustering Analysis

Hierarchical clustering was performed heatmap and dendrograms were performed utilising ‘gplots’, ‘dendextend’ and ‘colorspace’ packages in R with code adapted from [54,55], complete markdown can be found in **Supplementary Methods**. FPKM values were log10 transformed, and z-scored (Pearson Correlation method) prior to clustering. Hierarchical clustering resulted in distinct 6 clusters which are highlighted in different colours in the dendrogram (**Figure 4**)

#### Enrichment of association signal in IGAP GWAS data

As *Spi1* has been implicated in the regulation of other AD risk gene expression, both RNA-Seq datasets were tested for enrichment of association with AD risk in the International Genomics of Alzheimer’s Project (IGAP) GWAS dataset [56]. Firstly, the BioMart feature in Ensembl was used to convert mouse genes differentially expressed in the *Spi1* knock-down and over-expression datasets into human orthologs [57].

Gene sets were then determined for each *Spi1* RNA-Seq dataset using p-value cut-offs between 0.05 and 1×10^-10^. Direction of differential expression was not considered when defining gene sets. These gene sets (**Supplementary Table 2**) were tested for enrichment in the IGAP dataset using Multi-marker Analysis of GenoMic Annotation (MAGMA) analysis [58].

### Flow Cytometric Analysis of PU.1

Mixed glial cultures were harvested on day 21. Culture media was carefully removed and retained. Cells were detached by incubating with Accumax™ for 20 minutes at 37°C, added to the appropriate culture media and centrifuged at 300 *x g* for 7 minutes. Approximately 2×10^5^ cells per sample were fixed in 4 % formaldehyde solution (in DPBS) on ice for 30 minutes. Formaldehyde was removed by centrifugation (300 *x g* for 7 minutes) and cells were permeabilised on ice in 90 % ice cold methanol (v/v in PBS). Samples were centrifuged at 300 *x g* for 7 minutes and the supernatant discarded. To ensure the methanol was completely removed an additional wash step using 500 μL of wash solution (0.5 % (w/v) BSA, 5 mM EDTA and 2 mM NaN3 in DPBS) and centrifugation at 300 *x g* for 7 minutes. The supernatant was discarded, and the cell pellets were then re-suspended in 50 μL of block solution (5 % (v/v) filtered Rabbit Serum, 4 μg/mL Rat Anti-mouse FcγRII/III 2.4G2 clone in wash solution) and incubated on ice for 20-30 minutes. Following the blocking step 50 μL of CD11b, CD45 and PU.1 antibodies (listed below) were added and cells were incubated for 30-60 minutes on ice in the dark. The following antibodies were used in Flow cytometric experiments; Anti-CD11b FITC (56C, 8 μg/mL, in house), Anti-CD11b BV421 (M1/70, 2 μg/mL, Biolegend), Anti-CD11b PerCP-Cy5.5 (M1/70, 2 μg/mL, BD Biosciences), Anti-CD45 eFlour450 or PE-Cyanine7 conjugates (30-F11, 2 μg/mL, eBioscience), Anti-PU.1 AF647 (7C2C34, 5 μg/mL, Biolegend) and Rat IgG2a,k AF647 Isotype Control (RTK2758, 5 μg/mL, Biolegend).

Cells were washed with wash solution 3 times and centrifuged at 300 *x g* for 7 minutes. After the final wash the cell pellets were re-suspended in 500 μL of wash solution and acquired on Attune NxT cytometer (Thermofisher). Data was analysed using FlowJo software (version 10; FlowJo LLC). Single colour and isotype controls were used as appropriate. Post data collection relative PU.1 protein levels were determined using the Median Fluorescent Intensity (MFI) values in the PU.1 antibody channel. First the isotype background signal from the same channel was subtracted. These values were then normalised between experiments by diving each value by the average MFI value (minus isotype) of the appropriate control sample.

### Statistics and Figures

Statistical analyses were performed using GraphPad PRISM® 6 (version 3.07; GraphPad Software, Inc.), any statistical tests will be described as appropriate. P-values of >0.05 were taken as non-significant (ns). p-values of ≤0.05 will be denoted with a single asterisk*, p-values of ≤0.01 will be written as **, p-values of ≤0.001 by *** and p-value of ≤0.0001 as ****. Figures were made using both GraphPad PRISM® 6 (version 3.07; GraphPad Software, Inc.) and R Studio (Version 1.2.5042© 2009-2020 RStudio, Inc.) with base R version 4.0.0 (2020-04-24, Copyright © 2020 The R Foundation for Statistical Computing).

## Acknowledgements

We would like to thank our animal facility staff for the caring for the animals used in this study. We also thank the Cardiff University data clinic volunteers for their advice, we would like to especially thank Dr John Watkins for his guidance. We acknowledge Wales Gene Park and the Advanced Research Computing Team at Cardiff University, for performing Illumina® sequencing and provision of high-performance computing infrastructure. This project was funded by the MRC-Doctoral Training Programme Studentship. PRT is funded by a UK Dementia Research Institute Professorship and a Wellcome Trust Investigator Award (107964/Z/15/Z).

## Author contributions

RJ conducted the work; RA, PH and MH assisted with data analysis and interpretation; RJ and PRT conceived and designed the study and wrote the manuscript; All authors commented on and approved the manuscript.

## Additional Information

MH now works for Vertex Pharmaceuticals. The authors otherwise have no competing interests to declare.

## Notes

### Competing Interest Statement

M.H. is now an employee of Vertex Pharmaceuticals.

## References

1. Knapp PM, Guerchet M, McCrone M, Prina P, Comas-Herrera M, Wittenberg A, et al. Dementia UK Update. 2014.

2. Jones L, Holmans PA, Hamshere ML, Harold D, Moskvina V, Ivanov D, et al. Genetic Evidence Implicates the Immune System and Cholesterol Metabolism in the Aetiology of Alzheimer’s Disease. PLoS One. 2010 Nov 15;5(11):13950.

3. Escott-Price V, Bellenguez C, Wang LS, Choi SH, Harold D, Jones L, et al. Gene-wide analysis detects two new susceptibility genes for Alzheimer’s disease. PLoS One. 2014;9(6):94661.

4. Kunkle BW, Grenier-Boley B, Sims R, Bis JC, Damotte V, Naj AC, et al. Genetic meta-analysis of diagnosed Alzheimer’s disease identifies new risk loci and implicates Aβ, tau, immunity and lipid processing. Nat Genet. 2019;51:414–30.

5. Sims R, van der Lee SJ, Naj AC, Bellenguez C, Badarinarayan N, Jakobsdottir J, et al. Rare coding variants in PLCG2, ABI3, and TREM2 implicate microglial-mediated innate immunity in Alzheimer’s disease. Nat Genet. 2017;49(9):1373–84.

6. Gjoneska E, Pfenning AR, Mathys H, Quon G, Kundaje A, Tsai LH, et al. Conserved epigenomic signals in mice and humans reveal immune basis of Alzheimer’s disease. Nature. 2015;518:365–9.

7. Huang K, Marcora E, Pimenova AA, Narzo AF Di, Kapoor M, Jin SC, et al. A common haplotype lowers SPI1 (PU.1) expression in myeloid cells and delays age at onset for Alzheimer’s disease. Nat Neurosci. 2017;20:1052–61.

8. Tansey KE, Cameron D, Hill MJ. Genetic risk for Alzheimer’s disease is concentrated in specific macrophage and microglial transcriptional networks. Genome Med. 2018;10(14):1–10.

9. Celada A, Borras FE, Soler C, Lloberas J, Klemsz M, Beveren C van, et al. The Transcription Factor PU.1 Is Involved in Macrophage Proliferation. J Exp Med. 1996;184:61–9.

10. Anderson KL, Smith KA, Conners K, McKercher SR, Maki RA, Torbett BE. Myeloid Development Is Selectively Disrupted in PU.1 Null Mice. Blood. 1998 May 15;91(10):3702–10.

11. Beers DR, Henkel JS, Xiao Q, Zhao W, Wang J, Yen AA, et al. Wild-type microglia extend survival in PU.1 knockout mice with familial amyotrophic lateral sclerosis. PNAS. 2006 Oct 24;103(43):16021–6.

12. Scott EW, Simon MC, Anastasi J, Singh H. Requirement of transcription factor PU.1 in the development of multiple hematopoietic lineages. Science (80-). 1994 Sep;265:1573–7.

13. McKercher SR, Torbett BE, Anderson KL, Henkel GW, Vestal DJ, Baribault H, et al. Targeted disruption of the PU.1 gene results in multiple hematopoietic abnormalities. EMBO J. 1996;15(20):5647–58.

14. DeKoter RP, Singh H. Regulation of B lymphocyte and macrophage development by graded expression of PU.1. Science (80-). 2000 May;288:1439–41.

15. Back J, Dierich A, Bronn C, Kastner P, Chan S. PU.1 determines the self-renewal capacity of erythroid progenitor cells. Blood. 2004;103(10):3615–23.

16. Back J, Allman D, Chan S, Kastner P. Visualizing PU.1 activity during hematopoiesis. Exp Hematol. 2005;33:395–402.

17. Hu Z, Gu X, Baraoidan K, Ibanez V, Sharma A, Kadkol SH, et al. RUNX1 regulates corepressor interactions of PU.1. Blood. 2011;117(24):6498–508.

18. Jin H, Li L, Xu J, Zhen F, Zhu L, Liu PP, et al. Runx1 regulates embryonic myeloid fate choice in zebrafish through a negative feedback loop inhibiting Pu.1 expression. Blood. 2012;119(22):5239–49.

19. Zarnegar MA, Chen J, Rothenberg E V. Cell-Type-Specific Activation and Repression of PU.1 by a Complex of Discrete, Functionally Specialized cis-Regulatory Elements. Mol Cell Biol. 2010;30(20):4922–39.

20. Li Y, Okuno Y, Zhang P, Radomska HS, Chen H, Iwasaki H, et al. Regulation of the PU. 1 gene by distal elements. Blood. 2001;98(10):2958–65.

21. Okuno Y, Huang G, Rosenbauer F, Evans EK, Radomska HS, Iwasaki H, et al. Potential Autoregulation of Transcription Factor PU. 1 by an Upstream Regulatory Element. Mol Cell Biol. 2005;25(7):2832–45.

22. Zarnegar MA, Rothenberg E V. Ikaros represses and activates PU.1 cell-type-specifically through the multifunctional Sfpi1 URE and a myeloid specific enhancer. Oncogene. 2012 Oct 25;31:4647–54.

23. Leddin M, Perrod C, Hoogenkamp M, Ghani S, Assi S, Heinz S, et al. Two distinct auto-regulatory loops operate at the PU.1 locus in B cells and myeloid cells. Blood. 2011;117(10):2827–38.

24. Kamath MB, Houston IB, Janovski a J, Zhu X, Gowrisankar S, Jegga a G, et al. Dose-dependent repression of T-cell and natural killer cell genes by PU.1 enforces myeloid and B-cell identity. Leukemia. 2008;22:1214–25.

25. Lloberas J, Soler C, Celada A. The key role of PU.1/SPI-1 in B cells, myeloid cells and macrophages. Immunol Today. 1999 Apr;20(4):184–9.

26. Pahl HL, Scheibe RJ, Zhang DE, Chen HM, Galson DL, Maki RA, et al. The proto-oncogene PU.1 regulates expression of the myeloid-specific CD11b promoter. J Biol Chem. 1993;268(7):5014–20.

27. Anderson KL, Nelson SL, Perkin HB, Smith KA, Klemsz MJ, Torbett BE. PU.1 is a Lineage-specific Regulator of Tyrosine Phosphatase CD45. J Biol Chem. 2001;276(10):7637–42.

28. Heinz S, Benner C, Spann N, Bertolino E, Lin YC, Laslo P, et al. Simple Combinations of Lineage-Determining Transcription Factors Prime cis-Regulatory Elements Required for Macrophage and B Cell Identities. Mol Cell. 2010;38:576–89.

29. Ghisletti S, Barozzi I, Mietton F, Polletti S, De Santa F, Venturini E, et al. Identification and Characterization of Enhancers Controlling the Inflammatory Gene Expression Program in Macrophages. Immunity. 2010;32:317–28.

30. Gosselin D, Link VM, Romanoski CE, Fonseca GJ, Eichenfield DZ, Spann NJ, et al. Environment Drives Selection and Function of Enhancers Controlling Tissue-Specific Macrophage Identities. Cell. 2014 Dec;159:1327–40.

31. Rustenhoven J, Smith AM, Smyth LC, Jansson D, Scotter EL, Swanson ME V., et al. PU.1 regulates Alzheimer’s disease-associated genes in primary human microglia. Mol Neurodegener. 2018;13:44.

32. Pimenova AA, Herbinet M, Gupta I, Machlovi SI, Bowles KR, Marcora E, et al. Alzheimer’s-associated PU.1 expression levels regulate microglial inflammatory response. Neurobiol Dis. 2021;148:105217.

33. Olmos-Alonso A, Schetters STTT, Sri S, Askew K, Mancuso R, Vargas-Caballero M, et al. Pharmacological targeting of CSF1R inhibits microglial proliferation and prevents the progression of Alzheimer’s-like pathology. Brain. 2016 Jan 8;139:891–907.

34. Dagher NN, Najafi AR, Kayala KMN, Elmore MRP, White TE, Medeiros R, et al. Colony-stimulating factor 1 receptor inhibition prevents microglial plaque association and improves cognition in 3xTg-AD mice. J Neuroinflammation. 2015;12(139):1–14.

35. Sosna J, Philipp S, Albay RI, Reyes-Ruiz JM, Baglietto-Vargas D, LaFerla FM, et al. Early long-term administration of the CSF1R inhibitor PLX3397 ablates microglia and reduces accumulation of intraneuronal amyloid, neuritic plaque deposition and pre-fibrillar oligomers in 5XFAD mouse model of Alzheimer’s disease. Mol Neurodegener. 2018;13:1–11.

36. Spangenberg EE, Lee RJ, Najafi AR, Rice RA, Elmore MRP, Blurton-Jones M, et al. Eliminating microglia in Alzheimer’s mice prevents neuronal loss without modulating amyloid-β pathology. Brain. 2016;139:1265–81.

37. Smith AM, Gibbons HM, Oldfield RL, Bergin PM, Mee EW, Faull RLM, et al. The transcription factor PU.1 is critical for viability and function of human brain microglia. Glia. 2013 Jun 1;61:929–42.

38. Zhang DE, Hetherington CJ, Chen HM, Tenen DG. The macrophage transcription factor PU.1 directs tissue-specific expression of the macrophage colony-stimulating factor receptor. Mol Cell Biol. 1994;14(1):373–81.

39. Sierksma A, Lu A, Salta E, Mancuso R, Zoco J, Blum D, et al. Novel Alzheimer risk genes determine the microglia response to amyloid-β but not to TAU pathology. bioRxiv (preprint). 2019;491902.

40. Marecki S, Riendeau CJ, Liang MD, Fenton MJ. PU.1 and Multiple IFN Regulatory Factor Proteins Synergize to Mediate Transcriptional Activation of the Human IL-1β Gene. J Immunol. 2001;166:6892–6838.

41. Salih DA, Bayram S, Guelfi S, Reynolds RH, Shoai M, Ryten M, et al. Genetic variability in response to amyloid beta deposition influences Alzheimer’s disease risk. Brain Commun. 2019;1(1):1–13.

42. Schoggins JW. Interferon-Stimulated Genes: What Do They All Do? Annu Rev Virol. 2019 Sep 29;6(1):567–84.

43. Butovsky O, Jedrychowski MP, Moore CS, Cialic R, Lanser AJ, Gabriely G, et al. Identification of a unique TGF-β-dependent molecular and functional signature in microglia. Nat Neurosci. 2014 Jan;17(1):131–43.

44. Bohlen CJ, Bennett FC, Tucker AF, Collins HY, Mulinyawe SB, Barres BA. Diverse Requirements for Microglial Survival, Specification, and Function Revealed by Defined-Medium Cultures. Neuron. 2017;94:759–73.

45. Rosas M, Davies LC, Giles PJ, Liao C-T, Kharfan B, Stone TC, et al. The transcription factor Gata6 links tissue macrophage phenotype and proliferative renewal. Science (80-). 2014;344:645–8.

46. Ipseiz N, Czubala MA, Bart VMT, Davies LC, Jenkins RH, Brennan P, et al. Effective <em>In Vivo</em> Gene Modification in Mouse Tissue-Resident Peritoneal Macrophages by Intraperitoneal Delivery of Lentiviral Vectors. Mol Ther – Methods Clin Dev. 2020 Mar 13;16:21–31.

47. Bolger AM, Lohse M, Usadel B. Trimmomatic: a flexible trimmer for Illumina sequence data. Bioinformatics. 2014 Aug 1;30(15):2114–20.

48. Simon Andrews. FastQC A quality control tool for high throughput sequence data [Internet]. Babraham Bioinformatics. Available from: http://www.bioinformatics.babraham.ac.uk/projects/fastqc/

49. Dobin A, Davis CA, Schlesinger F, Drenkow J, Zaleski C, Jha S, et al. STAR: ultrafast universal RNA-seq aligner. Bioinformatics. 2013 Jan 1;29(1):15–21.

50. Liao Y, Smyth GK, Shi W. featureCounts: an efficient general purpose program for assigning sequence reads to genomic features. Bioinformatics. 2014 Apr 1;30(7):923–30.

51. Ensembl FTP Download [Internet]. Available from: ftp://ftp.ensembl.org/pub

52. Love MI, Huber W, Anders S. Moderated estimation of fold change and dispersion for RNA-seq data with DESeq2. Genome Biol. 2014 Dec;15(12):550.

53. Zheng R, Wan C, Mei S, Qin Q, Wu Q, Sun H, et al. Cistrome Data Browser: expanded datasets and new tools for gene regulatory analysis. Nucleic Acids Res. 2019 Jan 8;47(D1):D729–35.

54. Liu Y. How to Draw Heatmap with Colorful Dendrogram [Internet]. Yang’s Research Blog. 2018 [cited 2020 Apr 23]. p. 1. Available from: https://liuyanguu.github.io/post/2018/07/16/how-to-draw-heatmap-with-colorful-dendrogram/

55. Brandon Yeo. How to plot a Heatmap in Rstudio, the easy way [Internet]. 2019 [cited 2020 Apr 23]. Available from: https://www.youtube.com/watch?v=OWWHfXgRw3k

56. Lambert JC, Ibrahim-Verbaas CA, Harold D, Naj AC, Sims R, Bellenguez C, et al. Meta-analysis of 74,046 individuals identifies 11 new susceptibility loci for Alzheimer’s disease. Nat Genet. 2013;45(12):1452–8.

57. Herrero J. How to get all the orthologous genes between two species [Internet]. Ensembl Blog. 2009 [cited 2018 Nov 20]. Available from: http://www.ensembl.info/2009/01/21/how-to-get-all-the-orthologous-genes-between-two-species/

58. De Leeuw CA, Mooij JM, Heskes T, Posthuma D. MAGMA: Generalized Gene-Set Analysis of GWAS Data. PLoS Compuational Biol. 2015;11(4):1–19.

59. Soetaert K. Package ‘plot3D.’ cran R project; 2019.

