## Supplementary for "Modest changes in *Spi1* dosage reveal the potential for altered microglial function as seen in Alzheimer’s disease"

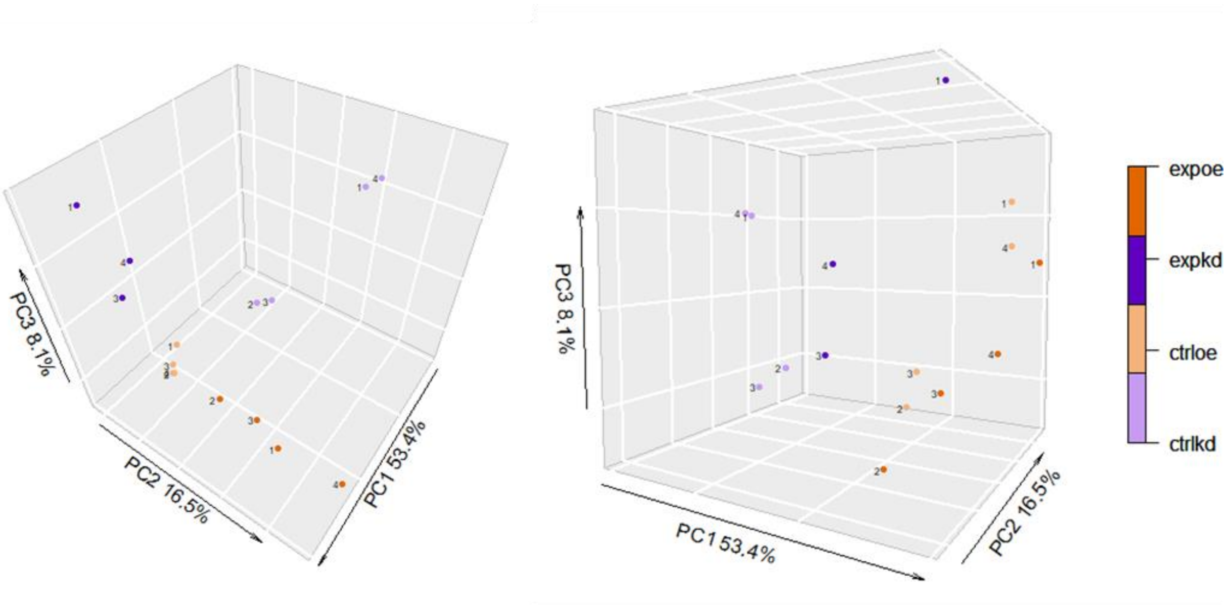


***Supplementary Figure 1- Plot Principal Components 1 and 2-*** *Each replicate within the sample group seems to have similar eigenvectors to the other samples (n=4 per group, except Spi1 shRNA where n=3). PC1 accounts for approximately half of the variation between samples, and it appears the control shRNA sample group is the most disparate. The top 10 genes that contribute to PC1 linked to processes such as mRNA processing/splicing and metabolic processes. The second and third principal components (PC2 & PC3) appears to represent the impact of Spi1 dose on the microglia transcriptome, as Spi1 pSIEW samples higher than the Spi1 shRNA samples, whereas both controls lie closer to the midline.*

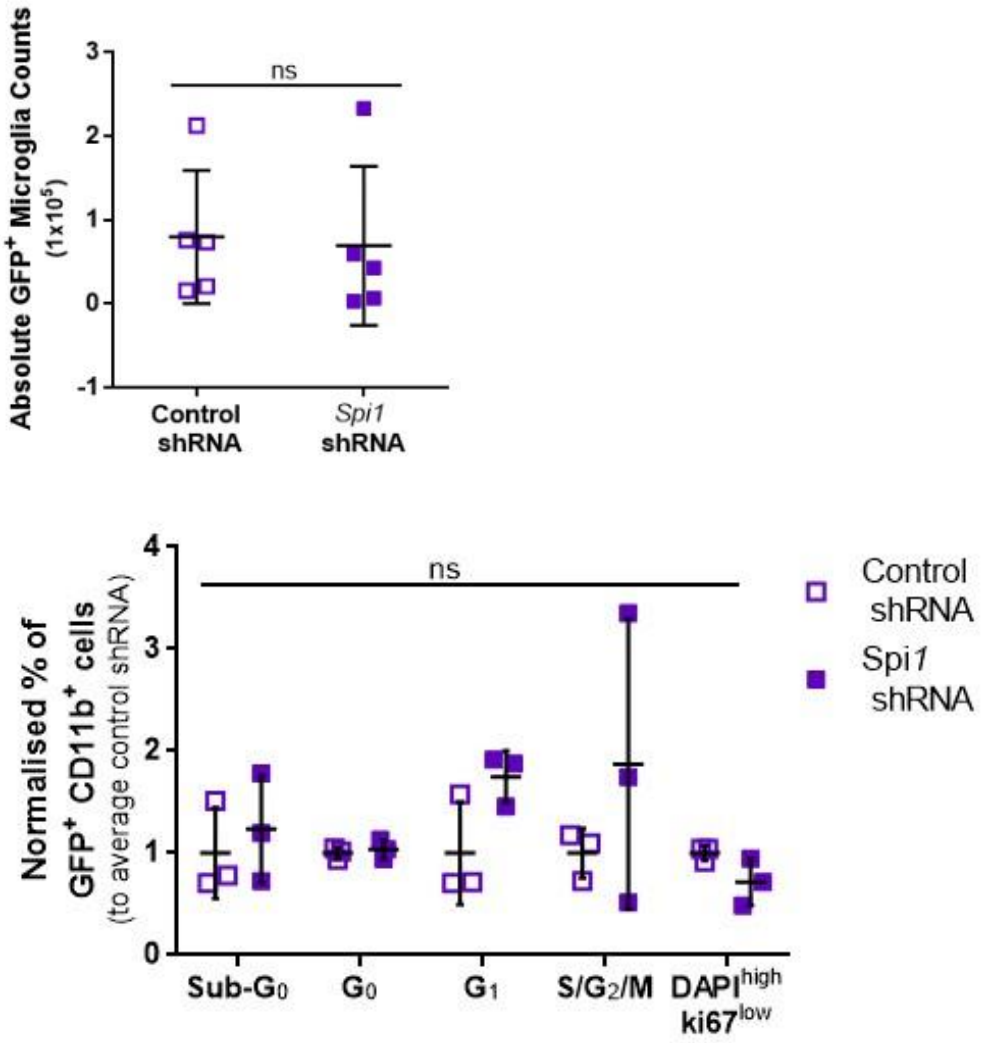


***Supplementary Figure 2- Functional validation of cell cycle alterations indicated by Spi1 knock-down RNA-Seq dataset. A*** *Absolute GFP+ microglia cell number did not differ between samples (One-tailed Paired t-Test, p-value = 0.1456, n=5).* ***B*** *The percentage of events in each cell cycle phase was normalised to the average proportion of control shRNA infected CD11b⁺ cells. Spi1 dose had a significant contribution to the variation in fold-change between groups it did not significantly impact the phases of the cell cycle (Two-Way ANOVA $P_{Spi1Dose} = 0.1202$, $P_{CellCycle} = 0.3406$, $P_{int} = 0.3406$).*

| Ensemble Gene ID | Gene Name | Ensemble Gene ID | Gene Name | Ensemble Gene ID | Gene Name |
|---|---|---|---|---|---|
| ENSMUSG00000000386 | Mx1 | ENSMUSG00000030126 | Tmcc1 | ENSMUSG00000046456 | Tmem150b |
| ENSMUSG00000000440 | Pparg | ENSMUSG00000030156 | Cd69 | ENSMUSG00000046879 | Irgm1 |
| ENSMUSG00000000552 | Zfp385a | ENSMUSG00000030275 | Etnk1 | ENSMUSG00000047712 | Ust |
| ENSMUSG00000001270 | Ckb | ENSMUSG00000030474 | Siglece | ENSMUSG00000052477 | C130026I21Rik |
| ENSMUSG00000002325 | Irf9 | ENSMUSG00000030683 | Sez6l2 | ENSMUSG00000052544 | St6galnac3 |
| ENSMUSG00000004562 | Arhgef40 | ENSMUSG00000030760 | Acer3 | ENSMUSG00000053063 | Clec12a |
| ENSMUSG00000006930 | Hap1 | ENSMUSG00000031785 | Adgrg1 | ENSMUSG00000053318 | Slamf8 |
| ENSMUSG00000009731 | Kcnd1 | ENSMUSG00000031838 | Ifi30 | ENSMUSG00000053559 | Smagp |
| ENSMUSG00000010358 | Ifi35 | ENSMUSG00000031839 | Hsbp1 | ENSMUSG00000053835 | H2-T24 |
| ENSMUSG00000016624 | Phf21b | ENSMUSG00000032420 | Nt5e | ENSMUSG00000053897 | Slc39a8 |
| ENSMUSG00000017830 | Dhx58 | ENSMUSG00000032640 | Chsy1 | ENSMUSG00000054404 | Slfn5 |
| ENSMUSG00000018341 | Il12rb2 | ENSMUSG00000032643 | Fhl3 | ENSMUSG00000055069 | Rab39 |
| ENSMUSG00000018459 | Slc13a3 | ENSMUSG00000032661 | Oas3 | ENSMUSG00000057219 | Armc7 |
| ENSMUSG00000019082 | Slc25a22 | ENSMUSG00000032690 | Oas2 | ENSMUSG00000057596 | Trim30d |
| ENSMUSG00000019843 | Fyn | ENSMUSG00000033355 | Rtp4 | ENSMUSG00000059895 | Ptp4a3 |
| ENSMUSG00000020023 | Tmcc3 | ENSMUSG00000033581 | Igf2bp2 | ENSMUSG00000060519 | Tor3a |
| ENSMUSG00000020261 | Slc36a1 | ENSMUSG00000034312 | Iqsec1 | ENSMUSG00000061132 | Blnk |
| ENSMUSG00000020638 | Cmpk2 | ENSMUSG00000034422 | Parp14 | ENSMUSG00000062488 | Ifit3b |
| ENSMUSG00000020641 | Rsad2 | ENSMUSG00000034438 | Gbp8 | ENSMUSG00000062960 | Kdr |
| ENSMUSG00000020788 | Atp2a3 | ENSMUSG00000034459 | Ifit1 | ENSMUSG00000063851 | Rnf183 |
| ENSMUSG00000020868 | Xylt2 | ENSMUSG00000034656 | Cacna1a | ENSMUSG00000066258 | Trim12a |
| ENSMUSG00000021087 | Rtn1 | ENSMUSG00000034783 | Cd207 | ENSMUSG00000066677 | Pydc3 |
| ENSMUSG00000021109 | Hif1a | ENSMUSG00000034842 | Art3 | ENSMUSG00000066861 | Oas1g |
| ENSMUSG00000021262 | Evl | ENSMUSG00000034855 | Cxcl10 | ENSMUSG00000068245 | Phf11d |
| ENSMUSG00000021263 | Degs2 | ENSMUSG00000035165 | Kcne3 | ENSMUSG00000069874 | Irgm2 |
| ENSMUSG00000021676 | Iqgap2 | ENSMUSG00000035208 | Slfn8 | ENSMUSG00000070034 | Sp110 |
| ENSMUSG00000021831 | Ero1l | ENSMUSG00000035352 | Ccl12 | ENSMUSG00000070327 | Rnf213 |
| ENSMUSG00000022756 | Slc7a4 | ENSMUSG00000035448 | Ccr3 | ENSMUSG00000070873 | Lilra5 |
| ENSMUSG00000022864 | D16Ertd472e | ENSMUSG00000035517 | Tdrd7 | ENSMUSG00000072889 | Nfxl1 |
| ENSMUSG00000022906 | Parp9 | ENSMUSG00000035692 | Isg15 | ENSMUSG00000073491 | Pydc4 |
| ENSMUSG00000023341 | Mx2 | ENSMUSG00000036986 | Pml | ENSMUSG00000073555 | Gm4951 |
| ENSMUSG00000023903 | Mmp25 | ENSMUSG00000037344 | Slc12a9 | ENSMUSG00000074896 | Ifit3 |
| ENSMUSG00000023988 | Bysl | ENSMUSG00000037406 | Htra4 | ENSMUSG00000075611 | Gm11545 |
| ENSMUSG00000024079 | Eif2ak2 | ENSMUSG00000037759 | Ptger2 | ENSMUSG00000076309 | Mir147 |
| ENSMUSG00000025165 | Sectm1a | ENSMUSG00000037816 | Fbxw17 | ENSMUSG00000076431 | Sox4 |
| ENSMUSG00000025279 | Dnase1l3 | ENSMUSG00000037849 | Gm4955 | ENSMUSG00000078452 | Raet1d |
| ENSMUSG00000025498 | Irf7 | ENSMUSG00000037921 | Ddx60 | ENSMUSG00000078616 | Trim30c |
| ENSMUSG00000025921 | Rdh10 | ENSMUSG00000037995 | Igsf9 | ENSMUSG00000078763 | Slfn1 |
| ENSMUSG00000026104 | Stat1 | ENSMUSG00000038068 | Rnf144b | ENSMUSG00000078853 | Igtp |
| ENSMUSG00000026222 | Sp100 | ENSMUSG00000038507 | Parp12 | ENSMUSG00000078920 | Ifi47 |
| ENSMUSG00000026536 | Mnda | ENSMUSG00000038518 | Jarid2 | ENSMUSG00000079363 | Gbp4 |
| ENSMUSG00000026790 | Odf2 | ENSMUSG00000038679 | Trps1 | ENSMUSG00000079484 | Phyhd1 |
| ENSMUSG00000026896 | Ifih1 | ENSMUSG00000038807 | Rap1gap2 | ENSMUSG00000082292 | Gm12250 |
| ENSMUSG00000026938 | Fcna | ENSMUSG00000038860 | Garnl3 | ENSMUSG00000084325 | Gm15550 |
| ENSMUSG00000027078 | Ube2l6 | ENSMUSG00000039236 | Isg20 | ENSMUSG00000085604 | Dhx58os |
| ENSMUSG00000027199 | Gatm | ENSMUSG00000039501 | Znfx1 | ENSMUSG00000085977 | Gm5970 |
| ENSMUSG00000027200 | Sema6d | ENSMUSG00000039899 | Fgl2 | ENSMUSG00000086481 | Gm11707 |
| ENSMUSG00000027368 | Dusp2 | ENSMUSG00000039967 | Zfp292 | ENSMUSG00000087700 | Gm15283 |
| ENSMUSG00000027514 | Zbp1 | ENSMUSG00000039976 | Tbc1d16 | ENSMUSG00000090125 | Pou3f1 |
| ENSMUSG00000028032 | Papss1 | ENSMUSG00000040033 | Stat2 | ENSMUSG00000090222 | Gm16340 |
| ENSMUSG00000028037 | Ifi44 | ENSMUSG00000040061 | Plcb2 | ENSMUSG00000091549 | Gm6548 |
| ENSMUSG00000028076 | Cd1d1 | ENSMUSG00000040296 | Ddx58 | ENSMUSG00000097461 | Gm26735 |
| ENSMUSG00000028268 | Gbp3 | ENSMUSG00000040328 | Olfr56 | ENSMUSG00000098871 | Mir6381 |
| ENSMUSG00000028270 | Gbp2 | ENSMUSG00000040365 | Trim41 | ENSMUSG00000101628 | Gm28177 |
| ENSMUSG00000028273 | Pdlim5 | ENSMUSG00000040483 | Xaf1 | ENSMUSG00000102715 | Gm6209 |
| ENSMUSG00000028378 | Ptgr1 | ENSMUSG00000040533 | Matn1 | ENSMUSG00000105504 | Gbp5 |
| ENSMUSG00000028514 | Usp24 | ENSMUSG00000041268 | Dmxl2 | ENSMUSG00000106586 | 1700013M08Rik |
| ENSMUSG00000028645 | Slc2a1 | ENSMUSG00000041827 | Oasl1 | ENSMUSG00000106734 | Gm20559 |
| ENSMUSG00000029298 | Gbp9 | ENSMUSG00000041992 | Rapgef5 | ENSMUSG00000107075 | Gm43068 |
| ENSMUSG00000029392 | Rilpl1 | ENSMUSG00000042106 | Fam212a | ENSMUSG00000107215 | Gm43197 |
| ENSMUSG00000029561 | Oasl2 | ENSMUSG00000042265 | Trem1 | ENSMUSG00000107222 | Gm43198 |
| ENSMUSG00000029605 | Oas1b | ENSMUSG00000044626 | Liph | ENSMUSG00000107736 | Gm44148 |
| ENSMUSG00000029798 | Herc6 | ENSMUSG00000044827 | Tlr1 | ENSMUSG00000107999 | Gm44123 |
| ENSMUSG00000030107 | Usp18 | ENSMUSG00000045932 | Ifit2 | ENSMUSG00000108112 | RP24-491O6.1 |
| | | ENSMUSG00000045973 | Slc25a51 | ENSMUSG00000109408 | RP23-25M3.6 |

***Supplementary Table 1- 194 Spi1 dose-sensitive genes including Ensemble Gene ID and Gene Name.***

| Experiment | RNA-Seq p-value Threshold | Number of genes | Enrichment Effect (β) | Enrichment p-value | Corrected Enrichment p-value |
|---|---|---|---|---|---|
| *Spi1* knock-down | 0.05 | 2380 | -0.01 | 0.7256 | 0.997 |
| | 0.01 | 1499 | -0.01 | 0.6272 | 0.992 |
| | 0.001 | 812 | -0.02 | 0.7864 | 0.998 |
| | 1.00E-04 | 476 | -0.03 | 0.7677 | 0.998 |
| | 1.00E-05 | 271 | -0.08 | 0.9444 | 1.000 |
| | 1.00E-06 | 166 | -0.04 | 0.7507 | 0.998 |
| | 1.00E-08 | 71 | 0.00 | 0.4971 | 0.972 |
| | 1.00E-10 | 33 | -0.02 | 0.5563 | 0.983 |
| *Spi1* over-expression | 0.05 | 267 | 0.12 | 0.0127 | 0.113 |
| | 0.01 | 145 | 0.10 | 0.0909 | 0.490 |
| | 0.001 | 78 | 0.15 | 0.0673 | 0.399 |
| | 1.00E-04 | 43 | 0.25 | 0.0356 | 0.253 |
| | 1.00E-05 | 28 | 0.42 | 0.0068 | 0.063 |
| | 1.00E-06 | 21 | 0.51 | 0.0033 | 0.036 |

***Supplementary Table 2- Information on the gene sets used in MAGMA analysis and the subsequent enrichment effects and p-values determined by this analysis.***

| Human_Entrez_ID | Human_Gene_Name | IGAP p-value | Mouse_Ensemble_ID | Mouse_Gene_Name | *Spi1* OE Fold_Change | *Spi1* OE adjusted p-value |
|---|---|---|---|---|---|---|
| 1949 | *EFNB3* | 5.64E-01 | ENSMUSG00000003934 | *Efnb3* | 5.3697 | 8.49E-24 |
| 93109 | *TMEM44* | 5.88E-01 | ENSMUSG00000022537 | *Tmem44* | 3.0809 | 1.61E-13 |
| 121551 | *BTBD11* | 4.95E-01 | ENSMUSG00000020042 | *Btbd11* | 3.0482 | 3.85E-12 |
| 167465 | *ZNF366* | 1.07E-02 | ENSMUSG00000050919 | *Zfp366* | 5.508 | 2.63E-11 |
| 93655 | *ST7* | N/A | ENSMUSG00000029534 | *St7* | 1.4258 | 7.47E-11 |
| 222236 | *NAPEPLD* | 2.84E-01 | ENSMUSG00000044968 | *Napepld* | 2.2926 | 2.36E-09 |
| 1152 | *CKB* | 8.34E-04 | ENSMUSG00000001270 | *Ckb* | 0.8262 | 6.26E-09 |
| 3437 | *IFIT3* | 3.07E-01 | ENSMUSG00000074896 | *Ifit3* | 2.3135 | 8.67E-09 |
| 730130 | *TMEM229A* | 4.66E-01 | ENSMUSG00000048022 | *Tmem229a* | 4.4963 | 8.67E-09 |
| 11269 | *DDX19B* | 7.11E-04 | ENSMUSG00000033658 | *Ddx19b* | 1.0481 | 1.93E-08 |
| 285852 | *TREML4* | 3.43E-03 | ENSMUSG00000051682 | *Treml4* | 2.3458 | 2.33E-08 |
| 4938 | *OAS1* | 3.24E-05 | ENSMUSG00000029605 | *Oas1b* | 1.5365 | 3.35E-08 |
| 6532 | *SLC6A4* | 4.14E-02 | ENSMUSG00000020838 | *Slc6a4* | 7.8789 | 4.08E-08 |
| 5732 | *PTGER2* | 5.02E-01 | ENSMUSG00000037759 | *Ptger2* | -1.2546 | 4.48E-08 |
| 23108 | *RAP1GAP2* | 1.24E-01 | ENSMUSG00000038807 | *Rap1gap2* | 1.055 | 4.71E-08 |
| 1776 | *DNASE1L3* | 6.73E-01 | ENSMUSG00000025279 | *Dnase1l3* | 4.837 | 1.00E-07 |
| 9636 | *ISG15* | 3.01E-01 | ENSMUSG00000035692 | *Isg15* | 1.9079 | 1.15E-07 |
| 255488 | *RNF144B* | 7.05E-01 | ENSMUSG00000038068 | *Rnf144b* | 3.9508 | 1.44E-07 |
| 55701 | *ARHGEF40* | 6.34E-01 | ENSMUSG00000004562 | *Arhgef40* | 0.4305 | 1.77E-07 |
| 4939 | *OAS2* | 5.25E-05 | ENSMUSG00000032690 | *Oas2* | 1.7721 | 1.84E-07 |
| 6252 | *RTN1* | 3.36E-01 | ENSMUSG00000021087 | *Rtn1* | 2.7493 | 2.10E-07 |
| 3750 | *KCND1* | N/A | ENSMUSG00000009731 | *Kcnd1* | 2.0755 | 4.49E-07 |
| 5175 | *PECAM1* | 7.25E-01 | ENSMUSG00000020717 | *Pecam1* | 3.0653 | 6.45E-07 |

***Supplementary Table 3- P-values for the 21 Spi1 over-expression RNA-Seq genes that were significantly enriched in the IGAP dataset.***

```
library(gplots)     # contains heatmap.2 function.

##
## Attaching package: 'gplots'

## The following object is masked from 'package:stats':
##
##     lowess

library(dendextend) # make and color dendrogram.

##
## ---------------------
## Welcome to dendextend version 1.13.4
## Type citation('dendextend') for how to cite the package.
##
## Type browseVignettes(package = 'dendextend') for the package vignette.
## The github page is: https://github.com/talgalili/dendextend/
##
## Suggestions and bug-reports can be submitted at: https://github.com/talgal
ili/dendextend/issues
## Or contact: <>
##
##  To suppress this message use:  suppressPackageStartupMessages(library(den
dextend))
## ---------------------

##
## Attaching package: 'dendextend'

## The following object is masked from 'package:stats':
##
##     cutree

library(colorspace)

#set working directory
setwd("C:/Users/rej66_000/OneDrive - Cardiff University/180626 RNA SEQ RESULT
S/For Clustering")

#Load FPKM values with p-value <=0.05 as csv.
df <- read.csv("FPKM.0.05.csv")

#renamed rows with gene ID then remove ID column.
row.names(df) <-df$geneID
df <- df[,-1]

#Convert the dataframe into a data matrix, required for analysis.
dm <- data.matrix(df)

#Scaling the data, Log10 scale and scale data so the mean is 0.
```

```r
Log_dm <- log10(dm+1)
SC_dm <- scale(Log_dm, scale = TRUE)
y <-SC_dm

#Hierarchical clustering of genes by row using pearson method, creating a den
drogram to show this.
dend1 <- as.dendrogram(hclust(as.dist(1-cor(t(y), method = "pearson")), metho
d = "complete"))
c_group <- 6                                              #number of cluste
rs data separated into.
dend1 <- color_branches(dend1, k = c_group, col = rainbow_hcl) #add color to
the clusters in the dendrograms.
dend1 <- reorder(dend1, rowMeans(y, na.rm = T), agglo.FUN = mean) #re-order t
he data by the dendrogram into clusters.

#Take the cluster colours from the dendrogram and apply to the ordered cluste
r label.
col_labels <- get_leaves_branches_col(dend1)
col_labels <- col_labels[order(order.dendrogram(dend1))]

#Colours for heatmap, where blue indicates low expression and red high expres
sion.
mycol <- colorpanel(75, "blue", "white", "red")

#Plot the heatmap with coloured dendrogram.
p <- heatmap.2(y, #scaled data matrix

               #Dendrogram and heatmap
               RowSideColors = col_labels,
               Rowv = dend1,
               Colv = FALSE,
               col = mycol,
               colRow = col_labels,
               density.info = "none",
               trace = "none",
               dendrogram = "row",

               #Key and labels
               key.title = "Scaled Z-Score",
               scale = "row",
               labRow = FALSE,
               cexCol = 1.2,
               labCol = c("NS shRNA 1", "NS shRNA 2","NS shRNA 3", "NS shRNA
4","Spi1 shRNA 1", "Spi1 shRNA 3", "Spi1 shRNA 4", "EV Control 1", "EV Contro
l 2", "EV Control 3", "EV Control 4", "Spi1 OE 1", "Spi1 OE 2", "Spi1 OE 3",
"Spi1 OE 4" ),
               offsetCol = -0.0000000001,
               margins = c(8,4),
               key = TRUE,
```

```
                #Creates a small space between sample sets
                colsep = c(4,7,11),
                sepcolor="white",
                sepwidth=c(0.05,0.05)
                 )
```

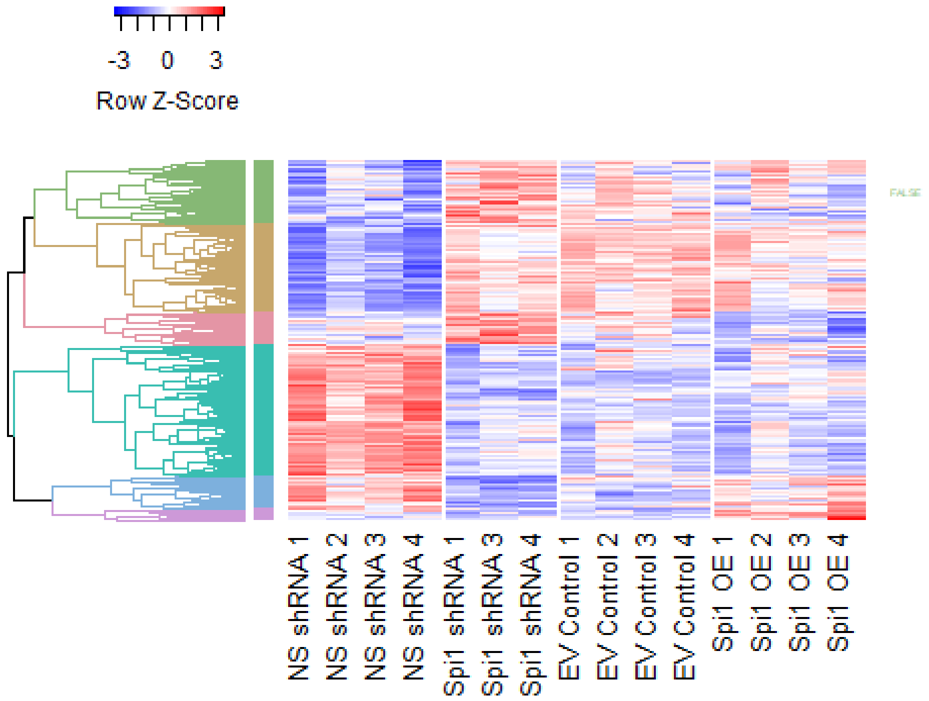


```
#Save cluster number against geneID
cluster <- as.matrix(col_labels)
output <- cbind(y, cluster)
write.csv(output, "R test Clustering output.csv", row.names = TRUE)

dev.off()

## null device
##           1
```

***Supplementary Figure 3 R Markdown for Clustering Analysis and generation of Figure 4a.***

### Supplementary Methods

#### Flow Cytometric Analysis of Ki67 Staining

Microglia were processed utilising a similar protocol that was flow cytometric analysis of PU.1, any differences are highlighted below. Briefly harvested cells were fixed in 1 % formaldehyde for 20 minutes on ice. Methanol permeabilization was not required as the wash solution was supplemented with 0.5 % Saponin. In addition to CD11b antibodies preciously mentioned, samples were stained with 2ug/mL of ki67 PE-Cyanine 7 (SolA15) or Rat IgG2a,k isotype control (eBR2a, both eBioscience™). Finally, 500ng/mL of 4',6-Diamidino-2-Phenylindole Dilactate (DAPI; Thermofisher) was added to each sample and incubated for 20 minutes before flow cytometric analysis.

Supplementary Figure 4 demonstrates the gating strategy utilised to analysis the separate cell cycle populations in infected microglia ($GFP^+CD11b^+$). PU.1 expression was confirmed to be reduced in *Spi1* shRNA infected samples compared to control shRNA infected microglia (data not shown).

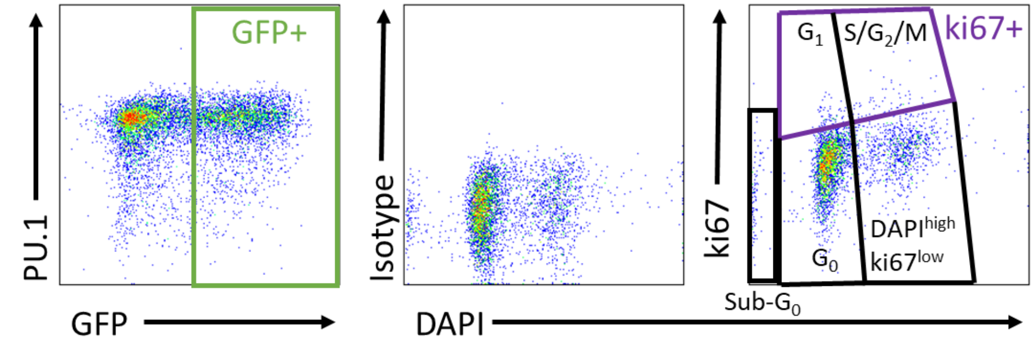


***Supplementary Figure 4 ki67 Gating Analysis-*** *$GFP^+$ microglia (determined as $CD11b^+$) were then divided into sub-populations of the cell cycle based on ki67 and DAPI staining as illustrated. In short, $ki67^+$ cells were defined as having a higher MFI than isotype controls which were then further separated into $G_1$ and $S/G_2/M$ based on DAPI staining. $G_0$, $DAPI^{high}$ cells were and sub-$G_0$ cells were defined as $ki67^-$ and separated according to the DAPI MFI.*

Absolute $GFP^+$ microglia number was calculated by dividing the percentage of $GFP^+$ microglia (of all events) by 100 and multiplying this by the total viable cell number, determined by Muse® Cell Analyzer (Merk Millipore) per manufacturer's directions.

#### Principle Component Analysis

Principle component analysis (PCA) was performed using the 'prcomp' function in R. A correlation matrix was generated from the FPKM normalised gene read counts reaching a significance threshold of $P \leq 0.05$, standardising the scores and giving them a mean of zero. The first three principle components (PC1, PC2 and PC3) accounted for 78 % of the variation between samples and were plotted using ‘plot3D’ package in R [59]. The loading scores for the top 10 genes contributing to PC1, PC2 and PC3 were also assessed.
